# Drug-driven reclassification of multiple tumour subtypes reveals intrinsic molecular concordance of therapy across histologically disparate cancers

**DOI:** 10.1101/2021.03.02.433516

**Authors:** Yue Xu, Jie Zheng, Zhaoqing Cai, Wang Li, Jens Köhler, Yao Dai, Xiaojie Cheng, Tao Wu, Fan Zhang, Haiyun Wang

**Affiliations:** School of Life Sciences and Technology, Tongji University, Shanghai 200092, China; Department of Thoracic Surgery, Shanghai Pulmonary Hospital, Tongji University School of Medicine, 507 Zhengmin Road, Shanghai 200433, China; Department of Medical Oncology, Dana-Farber Cancer Institute, Boston, MA 02215, USA

**Keywords:** Tumour classification, pharmacogenomics data, pharmacological subtypes, precision medicine, drug sensitivity

## Abstract

Cancers that are histologically defined as the same type of cancer often need a distinct therapy based on underlying heterogeneity; likewise, histologically disparate cancers can require similar treatment approaches due to intrinsic similarities. A comprehensive analysis integrated with drug response data and genomic alterations, particularly to reveal therapeutic concordance mechanisms across histologically disparate tumour subtypes, has not yet been fully exploited.

In this study, we used pharmacogenomic profiling data provided from the Cancer Genome Project (CGP) in a systematic in silico investigation of the pharmacological subtypes of cancers and the intrinsic concordance of molecular mechanisms leading to similar therapeutic responses across histologically disparate tumour subtypes. We further developed a novel approach to redefine cell-to-cell similarity and drug-to-drug similarity from the therapeutic concordance, providing a new point of view to study cancer heterogeneity.

Our study identified that histologically different tumours, such as malignant melanoma and colorectal adenocarcinoma, could belong to the same pharmacological subtype regarding drug sensitivity to MEK inhibitors, which was determined by their genomic alterations, high occurrence of BRAF or KRAS mutations. Therapeutic concordance for chemotherapy drugs was identified across histologically disparate hematological tumors mainly due to the extraordinary activation of the cell cycle in blood cancers. A subcluster of SCLC had a more similar profile with hematological tumors, and was associated with the malignant phenotype, with a higher level of MYC expression. We developed a website to store and visualize the pharmacological subtypes of drugs, as well as their connected genomic and expression alterations.

## Introduction

Traditional tumour classification based on histopathologic diagnosis and the TNM staging system offers highly practical guidance for surgical resection, regional radiotherapy or chemotherapies. However, due to complex intertumour and intratumour heterogeneity, cancer patients with such histological diagnoses often suffer from receiving effective drug treatment and therapeutic resistance [1–3]. One of the key mechanisms behind this significant challenge is the fact that cancers from the same tissue of origin often present quite different mechanisms of oncogenesis at the molecular level [2, 3]. Decades of studies have focused on finding molecular subtypes within histopathologically defined tumour types by analysing large-scale genomic, transcriptomic, proteomic and epigenomic alterations [4]. As a result, a variety of molecular signatures have been identified to distinguish intrinsic molecular subtypes associated with patient survival, prognosis and response to different therapeutic modalities. For example, BRAF mutation in melanoma [5]; EGFR-mutant lung adenocarcinomas [6]; luminal A, luminal B, HER2-enriched, basal-like and normal-like subtypes in breast cancer [7–9]; and four prominent genetic subtypes in Diffuse large B-cell lymphoma (DLBCL), termed MCD based on the co-occurrence of MYD88L265P and CD79B mutations, BN2 based on BCL6 fusions and NOTCH2 mutations, N1 based on NOTCH1 mutations, and EZB based on EZH2 mutations and BCL2 translocations [10] are several well-established molecular subtypes with explicit clinical significance.

With these advances in challenging cancer heterogeneity by identifying subtypes within the same tissues, emerging studies are uncovering facts in the other direction that cancers across disparate tissues of origin can explicitly share common molecular mechanism of oncogenesis [11, 12]. For example, one study based on a large-scale genomic analysis revealed that lung squamous, head and neck, and a subset of bladder cancers shared highly concordant signatures typified by TP53 alterations, TP63 amplifications, and high expression of immune and proliferation pathway genes, implicating that those different cancers perhaps require similar treatment approaches [12]. Another study based on large-scale genomic data found that TP53 and KRAS were mutually exclusive in COAD, READ, and LUAD, but significantly coexisted in PAAD. These observations reveal the feasibility of considering the same treatment strategy in different tumor types [13]. Similarly, HER2-targeted therapy may be applied to other cancer types analogous to breast cancer because ERBB2/HER2, which can be amplified in breast cancer, is also mutated and/or amplified in subsets of glioblastoma and gastric, serous endometrial, bladder and lung cancers [11]. In recent years, the FDA granted approval to larotrectinib, which had marked and durable antitumour activity in a variety of patients with RTK fusion-positive cancer, regardless of age or tumour tissue type [13]. Such examples explicitly illuminate a distinct avenue for reclassifying multiple tumour types independent of histopathologic diagnosis, highlighting that treatment approaches specifically discovered in one disease can be applied to another due to their intrinsic concordance of molecular patterns.

Currently, emerging efforts, such as pan-cancer analysis projects, are being conducted to comprehensively define commonalities and differences across cancer types and tissues of origin [11]. Nonetheless, integrative analysis, particularly towards revealing intrinsic therapeutic concordance across histologically disparate tumour subtypes, has not yet been fully exploited. Currently, a large-scale pharmacogenomics study, the Cancer Genome Project (CGP), provides high-throughput genomic information and pharmacological profiling of anticancer drugs across hundreds of cell lines that represent explicit molecular subtypes of histologically defined tumours [14]. Hence, we integrated drug response information to pharmacological reclassify tumour subtypes regardless of their tissues of origin. Tumour cells in the same class present similar drug responses, and those in different classes show varied drug responses. Furthermore, by integrating genomic alteration and expression information, we unravelled the intrinsic concordant molecular mechanism associated with the common drug response across histologically disparate cancers. Importantly, this research provides us with a purely therapy-oriented perspective to re-examine tumour classifications independent of histology subtypes.

## Material and Methods

Data from a large-scale pharmacogenomics study, the Cancer Genome Project (CGP), was accessible from its website: http://www.cancerrxgene.org. Gene expression, mutation and drug sensitivity data were downloaded. The CGP dataset includes 987 cell lines, genome-wide analysis of mutations, copy number variations and expression profiling, as well as the presence of commonly rearranged cancer genes, and 367 pharmacological profiles (dataset version 2020) [14]. In these data, the natural logarithm of the IC50 value represents the drug sensitivity value. IC50 is the half maximal inhibitory concentration of an anticancer drug, and a lower value means more sensitivity. Cell lines cannot be classified according to histological subtypes provided by TCGA, labelled as “UNCLASSIFIED”, were excluded from our study.

### Generating the pharmacological subtypes tree

The cell lines, IC50 of which are smaller than maximal tested concentrations, are defined as sensitive, otherwise resistant. Our method iteratively splits the cancer cells into two groups in a way that gains the best separation of drug sensitivity between two groups until reaching two terminal conditions: The p value of the difference of two groups’ drug sensitivity values is smaller than 0.05 or all cancer cells in a node are sensitive or resistant.

Suppose the set of all cells is *S_total_,* and each cell has its drug sensitivity value. The procedure to grow a pharmacological subtype tree is as follows:

1. Define the root node. Set *S* as the set of cancer cells in this node, and assign *S_total_* to *S.*
2. Define the root node as the current node.
3. *N* denotes the number of cells in the current node. If *N* is smaller than 6 or all cells in such node are sensitive or resistant, finish the procedure of growing the tree. Otherwise, a heuristic algorithm splits *S* into two child nodes, the left node and the right node, which achieves the greatest drug sensitivity difference.
4. If no way can be found to split the current node (*p* < 0.05), finish the procedure of growing the tree. Otherwise, the current node is split into two child nodes.
5. Define the left node and right node. Suppose the set of cells in the left node is *S_l_* and the set in the right node is *S_r_. S_l_* and *S_r_* are the subsets of *S.*
6. Define the left node as the current node. Set *S* as the set of cells in this node, and assign *S_l_* to *S.* Repeat steps 3-5.
7. Define the right node as the current node. Set *S* as the set of cells in this node, and assign *S_r_* to *S.* Repeat steps 3-5.

### Algorithm for splitting the node

We used a heuristic algorithm to search for a reasonable number of divisions to split *S* into *S_l_* and *S_r_.* Suppose the number of the cells set is *ns.* We first sorted the cells by IC50 ascending and divided the cells with higher drug sensitivity into the *S_l_* group and the cells with lower drug sensitivity into the *S_r_* group. Since each node at least 6 cells, a total of *ns* – 5 divisions were considered.

For each division, the Mann-Whitney U test was applied to calculate the *p* value of drug sensitivity between *S_l_* and *S_r_*. The division with the smallest *p* value was selected as the optimized one to split the cancer cells in the current node.

### The cell similarity and the drug similarity

The similarity of cells is calculated based on their response to the different drugs. For each drug *k,* a pharmacological subtypes tree *S^k^* is generated, consisting of *n* subtypes:

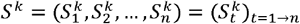

Based on this tree, a matrix that defines a similarity of cancer cells, regarding whether they are in the same subtype and their drug response to drug *k,* is calculated. We assume there are *m* cell lines tested for drug *k.* The cell lines are labelled as sensitive, resistant, or other ones. A sensitive cell line has its IC50 value smaller than minimum tested concentration; a resistant cell line has its IC50 value greater than maximal tested concentration. For two given cell lines *i* and *j,* their similarity matrix 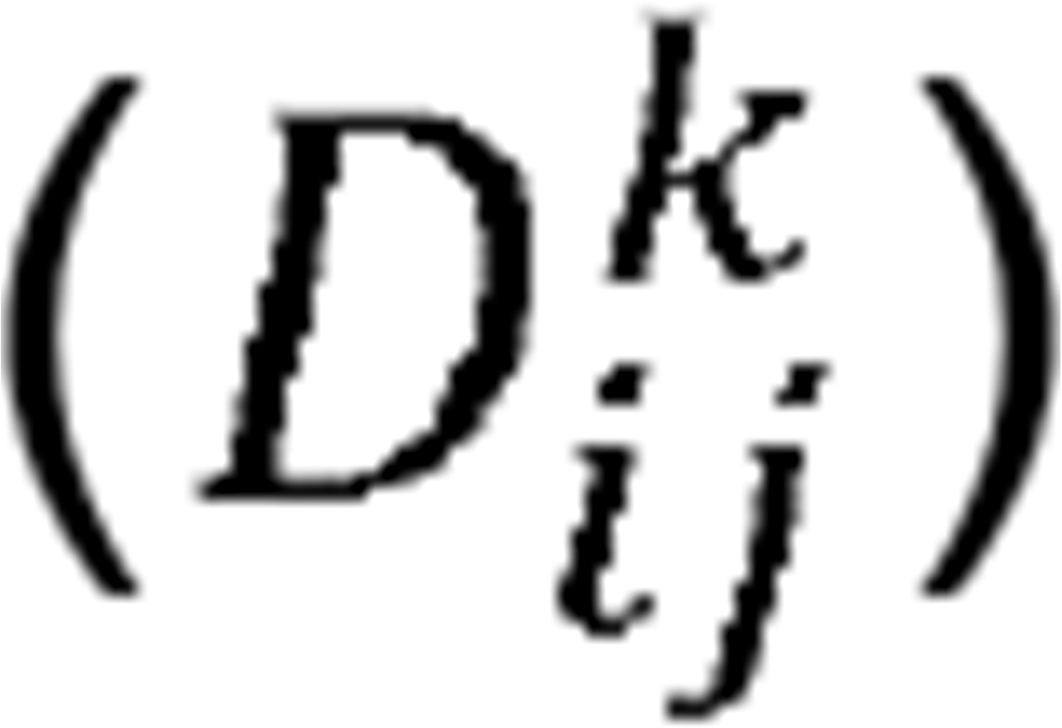 is defined as:

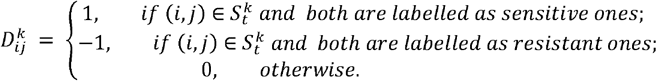

If two cell lines *i* and *j* are in the same subtype and both are sensitive to drug *k,* their score is 1, and −1 for both cell lines with a resistant response, while if they are not in the same subtype or show opposite response to the drug, their score is 0. A *m×m* similarity matrix of cell lines is then generated.

Then, the similarity of cell lines i and *j* can be further calculated as:

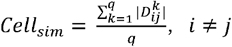

Wherein *i* and *j* denote the cell lines tested for *q* drugs.

Furthermore, for two given drugs *a* and *b*, their similarity is calculated as the summarized similarity across the same cell lines which are tested for both drugs.

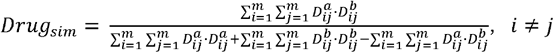

Wherein *i* and *j* denote the cell lines tested for both drugs, and *m* is the total number of the same cell lines.

### Connecting pharmacological subtypes with genomic alterations

The chi-squared test was used to calculate the connection between genomic alterations, including mutation and translocation, and pharmacological subtypes. Genomic alterations were determined with *p* values corrected with the Benjamini-Hochberg method for controlling the false discovery rate [15]. Here we used the corrected *p* value < 0.05 as the threshold. We defined two types of connections: positive and negative. If pharmacological subtypes with high drug sensitivity had more frequent alteration occurrences, we defined this connection as positive. If the opposite was true, we defined the connection as negative.

The connection between mRNA expression and pharmacological subtypes was determined with *p* values from the Kruskal-Wallis H test corrected with the Benjamini-Hochberg method. If pharmacological subtypes with high drug sensitivity had higher expression, we defined this connection as positive. If the opposite was true, we defined the connection as negative. Here we used the corrected *p* value < 0.05 as the threshold. The individual subtypes with sensitive and resistant cells mixed were excluded in the above analyses.

### Gene Set Enrichment Analysis (GSEA)

Gene set enrichment analysis (GSEA) [16] was employed to determine the 333 gene sets from KEGG, enriched by a pre-ranked list of all genes, which were sorted by the statistical significance of differential expression defined by DESeq2 analysis [17]. Gene sets with FDR < 0.05 were statistically significant.

### Statistical analysis

The Fisher’s exact test was used to respectively determine whether there is a significant association between pathways and clusters of drugs determined by the number of drug pharmacological subtypes across 6 cancer types or not, whether any difference of phenotypes (cancers/pathways) in clusters of cell lines/drugs derived from the similarity matrix was significant, and whether cancer censor genes are enriched in a list of genes with the most connections to pharmacological subtypes. A hypothesis test based on hypergeometric distribution was used to determine whether a histological cancer is enriched in the most sensitive pharmacological subtype of a drug. The *p* value < 0.05 was regarded as statistical significant. Kolmogorov-Smirnov test (K-S test) was employed to compare cumulative distribution function (CDF) of the number of pharmacological subtypes across the drugs between two histological cancers.

## Results

We analysed 367 drugs in the CGP dataset and established their pharmacological subtype trees. The leaf node in each tree represents a pharmacological subtype of cancers, which is composed of cells from the histological disparate tumour subtypes. We further employed the mRNA expression profile and mutation/fusion profile to connect the molecular alterations with pharmacological subtypes, delineating a novel perspective to re-classify the tumour according to therapeutic response dependent of the intrinsic concordant molecular mechanism, by regardless of tissues of origin. The cell similarity and the drug similarity were also redefined based on the pharmacological subtypes.

### Pharmacological subtypes of cancers

We built a pharmacological tree for each drug based on the divisibility of the drug sensitivity of cancer cells. Taking the tree of the MEK1/2 inhibitor PD0325901 as an example (Fig. 1A), all cells in the root node were first divided into left and right child nodes with relatively high and low sensitivity, respectively, and further cells in these two nodes were capable of being divided into six final subgroups, C1, C2, C3, C4, C5, and C6. Each subgroup had varied drug sensitivity and reached maximum divisibility, thus representing distinct pharmacological subtypes. C1 denotes the most sensitive and C6 the most resistant subtype to the drug PD0325901. Except for C4 where both sensitive and resistant cells were mixed, the other subgroups contained the homologous sensitive or resistant cells (Fig. 1B). There were much more sensitive cells in C1, C2 and C3 than resistant cells in C5 and C6 (Fig. 1C). We then investigated how histological subtypes were distributed in the pharmacological subtypes (Fig. 1D). C1 was composed of 22 histological subtypes, with top 2 cancers, SKCM (19.6%) and COREAD (11.3%). C6 was composed of 15 histological subtypes, with leading cancers including SCLC (36.6%) and BRCA (17.1%).

**Figure 1.**
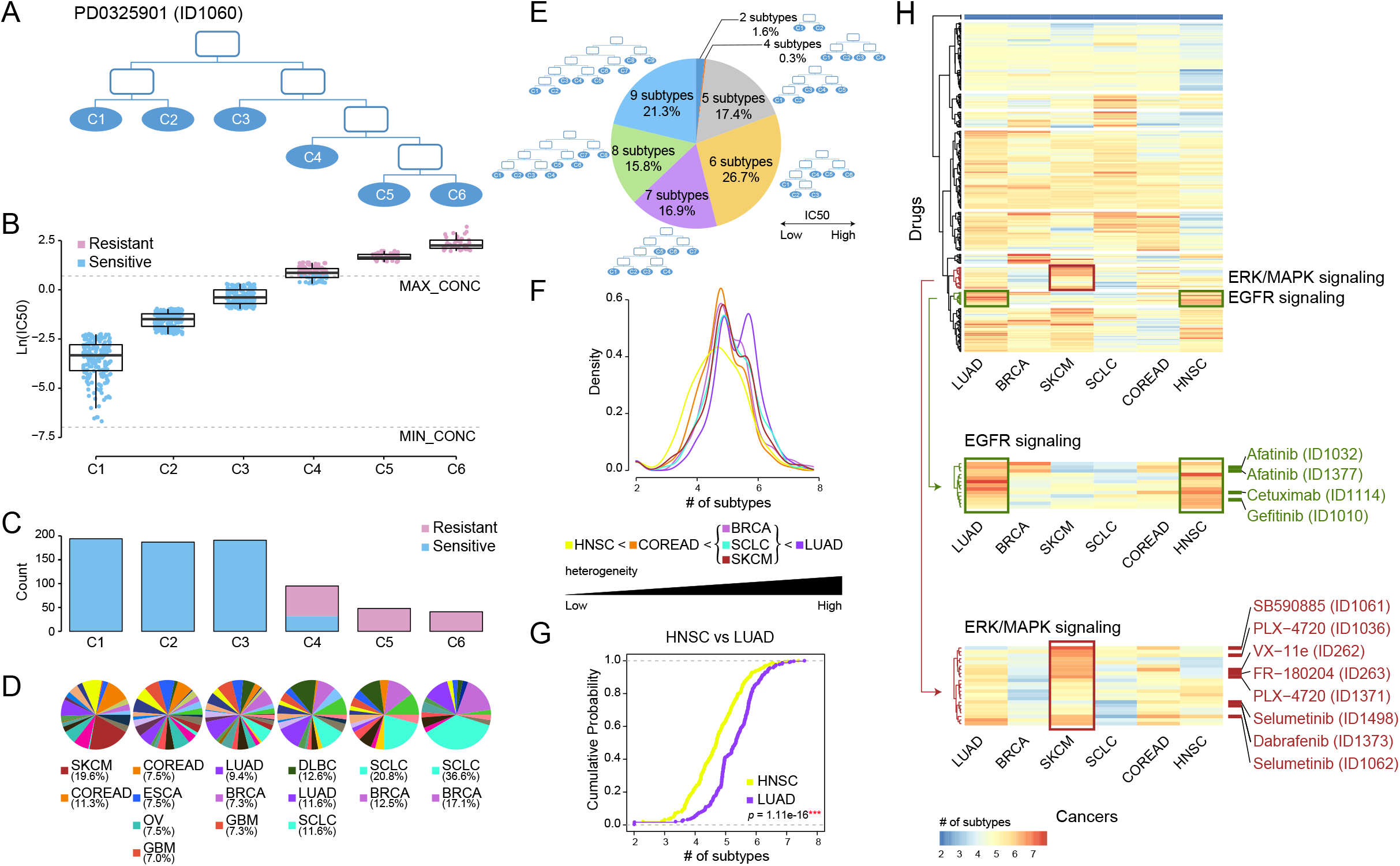
Re-classifying cancers to generate pharmacological subtypes and drug sensitivity profiles. (A) Pharmacological subtype tree of the drug PD0325901. Cancer cells were classified into six pharmacological subtypes based on their drug sensitivity to MEK inhibitor PD0325901. (B) The boxplot shows the drug sensitivities of six subtypes (leaf nodes): C1, C2, C3, C4, C5, and C6, indicating a gradually increasing degree of resistance to the drug PD0325901. Blue points refer to cell lines sensitive to the drug, while red points resistant. (C) The barplot shows the number of cell lines in each subtype. (D) The pie chart shows the distribution of histological subtypes across pharmacological subtypes. (E) The pie chart illustrates the numerical proportion of different degrees of the furthest divisible pharmacological subtypes. Cancer cells could be maximally classified into up to 9 subtypes for different drugs. Among them, 7-9 subtypes accounted for more than half of drugs. (F) The density of the number of pharmacological subtypes indicates the varied treatment heterogeneity across six histological cancers, in which the average number of cell lines among all drugs are over 35. (G) The empirical cumulative distribution function visualizes the density of drug subtypes in LUAD vs. HNSC. P value was calculated by using the Kolmogorov-Smirnov test (K-S test). (H) Hierarchical clustering of the number of pharmacological subtypes across six cancers. Each row in the heatmap represents a drug, and each cell in the heatmap represents the number of subtypes of the cancer in the drug. The red colour indicates more subtypes, and the blue colour less subtypes.

Our analysis established 367 pharmacological trees. The number of pharmacological subtypes with distinct drug sensitivities varied from 2 to 9 across 367 trees (Fig. 1E). This suggested that heterogeneity of treatment effects across histological subtypes existed widely, and one pharmacological subtype presenting concordant drug response indeed comprised histological disparate cells to some extent. A high degree of treatment heterogeneity, meaning more than seven pharmacological subtypes, was observed for 54.0% (16.9% of the drugs had 7; 15.8% of the drugs had 8; and 21.3% of the drugs had 9 pharmacological subtypes) of the drugs. These drugs included Dabrafenib (BRAF inhibitor), Selumetinib (MEK1/2 inhibitor), Erlotinib (EGFR inhibitor), and Alectinib (ALK inhibitor). 44.1% of the drugs had 5 or 6 subtypes, showing a moderate degree of treatment heterogeneity, while 1.9% of the drugs had subtypes lower than 4, showing a low degree of heterogeneity related to drug therapy (Fig. 1E).

We also particularly examined how pharmacological subtypes were constituted within the same histological tumour subtypes, including LUAD, BRCA, SKCM, SCLC, COREAD and HNSC, in which the average number of cell lines per drug was more than 35 (Fig. 1F-H). We built approximately 310 pharmacological trees for each type of the above cancers. The density distribution of the number of pharmacological subtypes indicated the varied treatment heterogeneity across 6 histological subtypes (Fig. 1F). The shape of the distribution curve in LUAD was characterized by double kurtosis and significantly positively biased, and oppositely the curve in HNSC was negatively biased, suggesting the highest treatment heterogeneity in LUAD and lowest treatment heterogeneity in HNSC. Furthermore, K-S test was applied to evaluate if two different histological subtypes had the same level of treatment heterogeneity by comparing their cumulative distributions. The results showed the cumulative distribution curve of LUAD is significantly different from that of HNSC (*p=1.11e-16,* K-S test) (Fig. 1G). The pairwise comparisons between the curves of any two histological subtypes showed BRCA, SCLC, and SKCM had the statistically similar distributions (*p>0.05,* K-S test), and other cancers had not (*p<0.05,* K-S test) (Supple. Fig. 1).

To observe the varied treatment heterogeneity of the different histological subtypes in detail, the number of pharmacological subtypes of drugs across 6 histological cancers were shown in the heatmap (Fig. 1H). Six histological tumors were arranged in columns according to their treatment heterogeneity, from high to low. Interestingly we observed that EGFR inhibitors (Afatinib, Cetuximab and Gefitinib) targeting in EGFR signalling pathway (green drugs in Fig. 1H) were clustered together (*p*<0.001, Fisher’s exact test). And in LUAD and HNSC there were more pharmacological subtypes of EGFR inhibitors than other drugs, suggesting high treatment heterogeneity for EGFR inhibitors in LUAD and HNSC. In addition, drugs targeting ERK/MAPK signalling (brown drugs in Fig. 1H) were clustered together (*p*<0.001, Fisher’s exact test). There was high treatment heterogeneity for these drugs in SKCM.

### Genomic alterations associated with pharmacological subtypes

To identify the molecular alterations associated with varied drug sensitivity, we examined whether the genomic alterations, including mutations and translocations, significantly changed across the pharmacological subtypes. The statistical significance was determined by *p* values calculated with the Chi-squared test and corrected with the Benjamini-Hochberg method for controlling the false discovery rate. If the pharmacological subtypes with higher drug sensitivity had more frequent mutation or translocation events, we defined their association as a positive connection. If the opposite was true, we defined their association as negative. Consequently, genes with specific genomic alterations were associated with the respective drugs. Since one gene could have multiple connected drugs and one drug could have multiple connected genes, these gene-drug connections ultimately composed a network that allowed us to investigate the contribution of either a drug or gene to the holistic understanding of how altered genes connect to drug responses (Fig. 2A).

**Figure 2.**
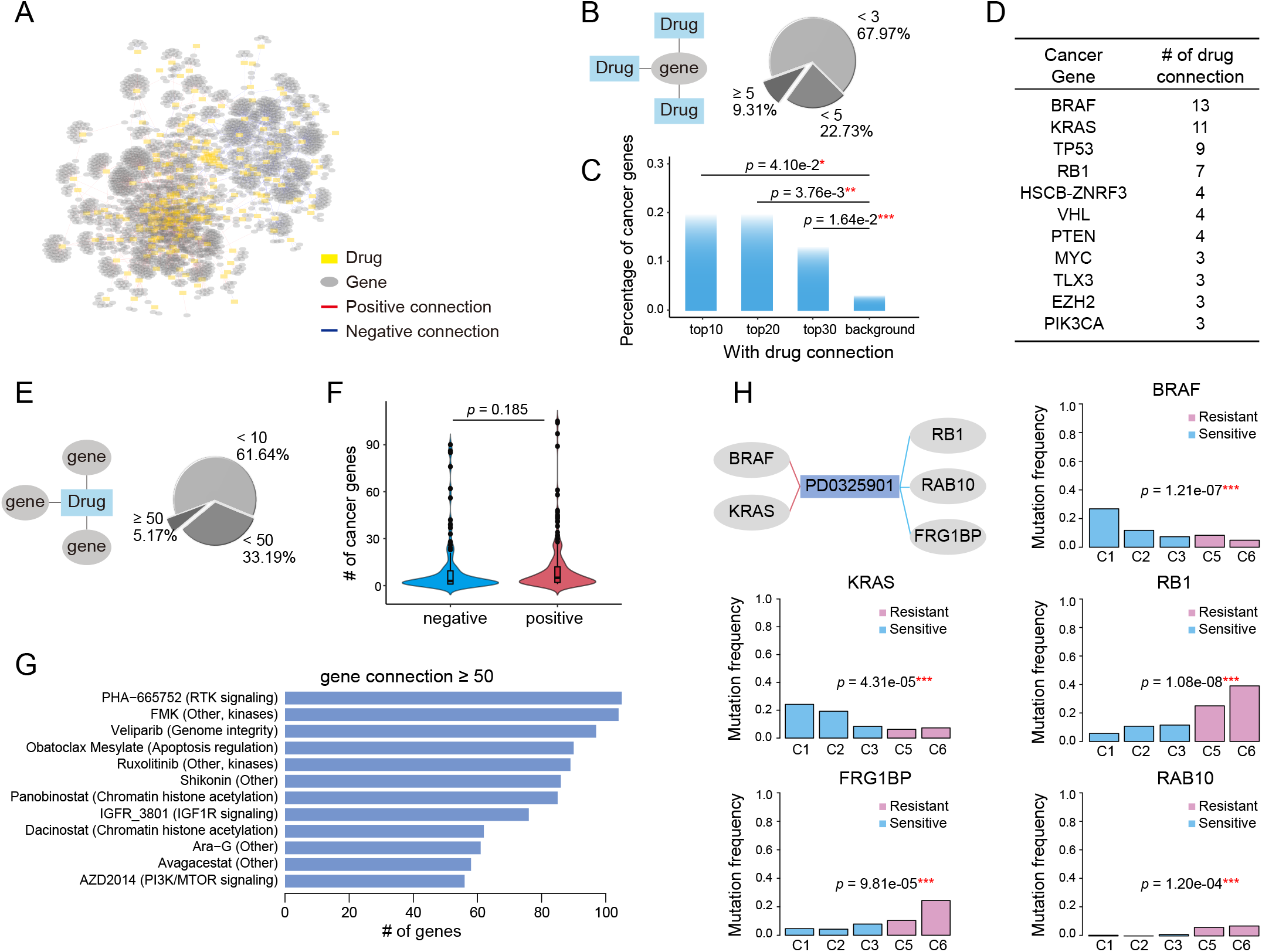
Genomic alterations determine the various drug sensitivities across pharmacological subtypes. (A) A gene with its genomic alteration significantly changed across the pharmacological subtypes of a drug is linked with the drug, generating a gene-drug network. A positive connection meant that subtypes with high drug sensitivity had a more frequent occurrence of genomic alterations. Conversely, those that were less frequent were defined as negative connections. (B) 9.31% genes had more than 5 connected drugs. In addition, 67.97% of genes had fewer than 3 connected drugs. (C) The percentage of cancer genes in the top 10, 20, and 30 genes, and the genome-wide 21,972 genes were referred to as background. (D) The top cancer censor genes defined by COSMIC and the corresponding number of drug connections. (E) When considering the genes connected to the drugs, 5.17% of drugs have more than 50 connected cancer genes. In addition, 61.64% of drugs had fewer than 10 connected cancer genes. (F) Comparison of the number of genes with a positive connection with the number of genes with a negative connection. (G) Drugs with cancer gene connections greater than 50 were ranked. (H) BRAF mutation, RB1 mutation, KRAS mutation, FRG1BP mutation and RAB10 mutation were associated with the five pharmacological subtypes of PD0325901. Histograms show the mutation/translocation ratio of the five genes in the five pharmacological subtypes of PD0325901.

We first investigated the distribution of genes on their connected drugs (Fig. 2B). Our analysis identified 462 genes whose genomic alterations were connected with at least two drugs’ pharmacological subtypes. Most of the genes (67.97%) had fewer than 3 connections with drugs, but 43 (9.31%) genes whose genomic alterations were associated with more than 5 drugs. By ranking genes by their number of connections to drugs, we found that the top ranked genes were enriched in cancer genes (Fig. 2C; Supple. Fig. 2), a catalogue of genes with mutations that are causally implicated in cancer provided by the COSMIC database [18]. The top 10 genes connected to more than 9 drugs, 20% of which were remarkably annotated as cancer genes (Fig. 2C). These cancerous percentages for the top 10 (*p*=4.10e-2, Fisher’s exact test), top 20 (*p*=3.76e-3, Fisher’s exact test), and top 30 genes (*p*=1.64e-2, Fisher’s exact test) were significantly higher than the background when using genome-wide 21,972 genes as a reference (Fig. 2C). Among the top ranked oncogenes, BRAF ranked first with connections to 13 drugs, followed by KRAS and TP53 with 11 and 9 drugs (Fig. 2D), implying that the highly drug-connected genes that are not currently identified as cancer genes could actually be causally implicated in cancer therapy, such as ADK-VCL fusion, TNFRSF9 and LTB, which showed connections to over 20 drugs and ranked as the top five genes (Supple. Fig. 2).

We then investigated the distribution of drugs on their connected genes. For the majority of drugs (61.64%), the number of connected genes ranged from 2 to 10 (Fig. 2E). There was the same number of genes that showed a positive connection as that showed a negative connection (Fig. 2F). We ranked the drugs with more than 50 genes’ connection with their pharmacological subtypes (Fig. 2G). Drugs with gene connections greater than 50 included PHA-665752, affecting the RTK signaling pathway, Veliparib, affecting the genome integrity pathway, Obatoclax Mesylate, affecting the apoptosis regulation pathway, and AZD2014, affecting the PI3K/MTOR signaling pathway.

Next, we observed how the mutation rate of genes associated with the variability of drug sensitivity changed among drug-sensitive and drug-resistant subtypes. Taking the MEK1/2 inhibitor PD0325901 as an example, five genomic alterations were found to connect to the drug and be associated with pharmacological subtypes. As shown in Fig. 2H, BRAF mutations and KRAS mutations occurred significantly more frequently (over 20%) in the sensitive subtype C1 and then decreased gradually in the subtypes C2, C3, C5 and C6. Here one subtype C4 with sensitive and resistant cells mixed was excluded in the analysis. Conversely, RB1 mutation, FRG1BP mutation and RAB10 mutation occurred more frequently in the resistant subtypes. Therefore, C1 group, consisting mainly of SKCM (19.6%) and COREAD (Fig. 1D), was significantly sensitive to PD0325901 due to its high occurrence of BRAF and KRAS mutations and low occurrence of RB1, FRG1BP and RAB10 mutations (Fig. 2H). Other types of MEK inhibitors, including Refametinib, Trametinib, Selumetinib, and CI-1040, further confirmed that BRAF, KRAS, and RB1 mutations were robustly connected to pharmacological subtypes, contributing to the drug sensitivity of MEK inhibitors (Supple. Fig. 3). Our analysis revealed that the varied distribution of genomic alterations across pharmacological subtypes could lead to their differences in response to anticancer therapies.

### Expression alterations associated with pharmacological subtypes

We also associated mRNA expression with the pharmacological subtypes. The connection between gene expression and pharmacological subtypes was determined by p values calculated with the Kruskal-Wallis H test and corrected with the Benjamini-Hochberg method for controlling the false discovery rate. If the subtypes with high drug sensitivity had higher gene expression, we defined this connection as positive. If the opposite was true, we defined the connection as negative. The connections between genes and drugs constituted a network (Fig. 3A). Since some genes whose expression was connected to pharmacological subtypes were indirectly associated with drug sensitivity, we then removed genes that had functions outside of the core cancer pathways (Supple. Tab. 1).

**Figure 3.**
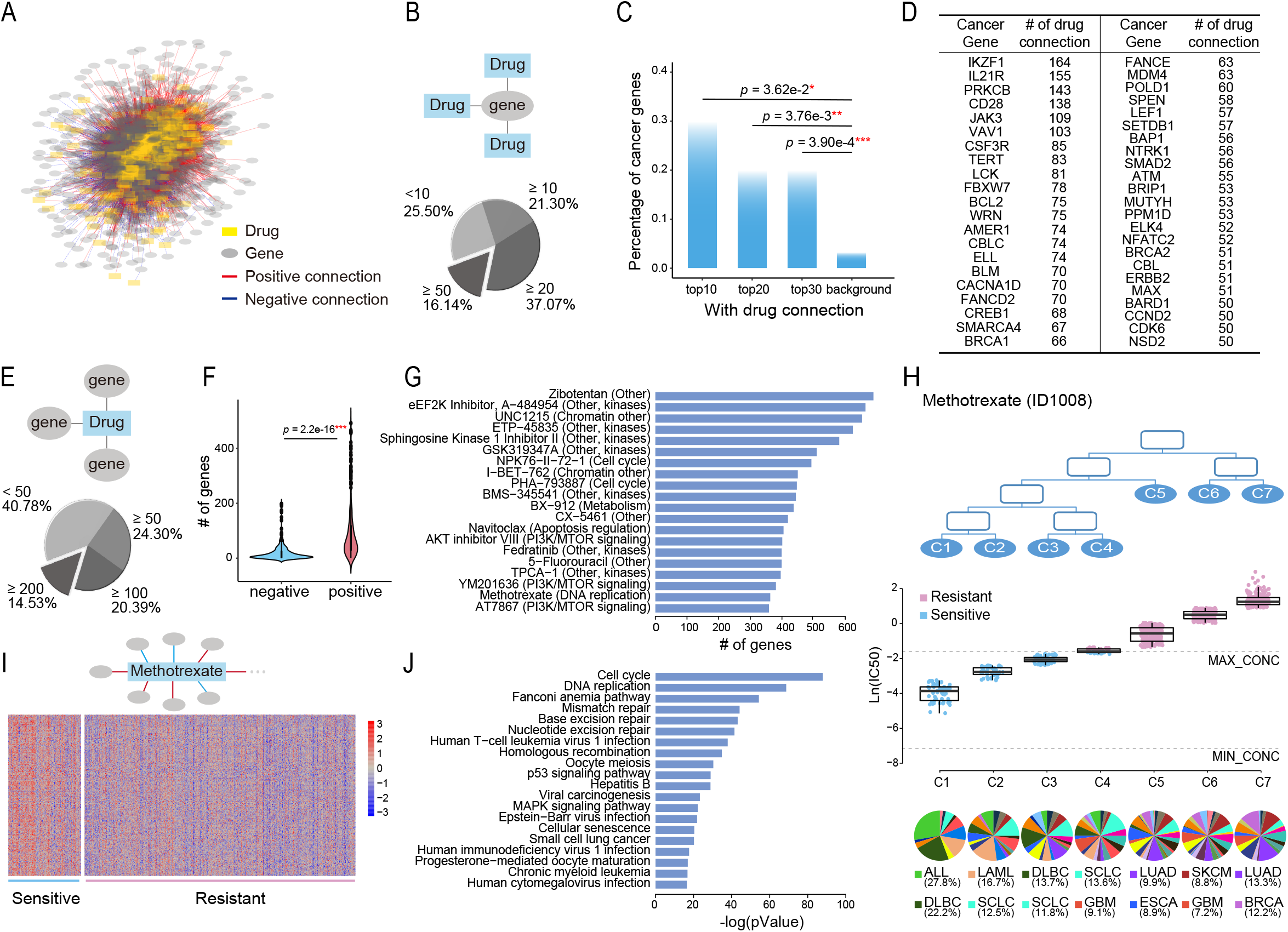
Tissue-specific gene expression determines the various drug sensitivities across pharmacological subtypes. (A) A gene with its expression significantly changed across the pharmacological subtypes of a drug is linked with this drug, generating a gene-drug network. A positive connection meant that subtypes with high drug sensitivity had higher gene expression. Conversely, those with low expression were defined as negative connections. (B) Pie chart illustrating the numerical proportion of the number of connected drugs with genes whose expression was associated with the pharmacological subtypes. A total of 16.14% of genes had more than 50 connected drugs. In addition, 25.50% of genes had less than 10 connected drugs. (C) The percentage of cancer genes in the top 10, 20, and 30 genes, and the total cancer genes were referred to as background. (D) The top cancer censor genes defined by COSMIC and the corresponding number of drug connections. (E) Distribution of the genes functioning in the core cancer pathways whose expression was connected with pharmacological subtypes. A total of 14.53% of drugs had more than 200 connected genes. (F) The number of genes with positive connections was significantly greater than the number of genes with negative connections. (G) Top 20 drugs with the most connections to genes. (H) Seven subtypes, C1, C2, C3, C4, C5, C6 and C7, had varied drug sensitivity to methotrexate, with C1 being the most sensitive to the drug and C7 being the most resistant to the drug. The boxplot below shows the drug sensitivities of the three subtypes (leaf nodes) C1, C2, C3, C4, C5, C6 and C7, indicating a gradually increasing degree of resistance to the drug. (I) A total of 297 genes were highly expressed in sensitive group (C1, C2 and C3) and lowly expressed in resistant group (C5, C6 and C7). (J) Pathways enriched by the 297 highly expressed genes in sensitive group (C1, C2 and C3).

We first investigated the distribution of genes on their connected drugs (Fig. 3B). Our analysis identified 1,357 genes whose expression was connected to at least two drugs’ pharmacological subtype. Among them, 16.14%, 37.07% and 21.30% of genes respectively had connections with greater than 50, 20 and 10 drugs. The top 10 genes connected to more than 143 drugs, 30% of which were annotated as cancer genes (Fig. 3C; Supple. Fig. 4). These cancerous percentages for the top 10 (*p*=3.62e-2, Fisher’s exact test), top 20 (*p*=3.67e-3, Fisher’s exact test), and top 30 genes (*p*=3.90e-4, Fisher’s exact test) were significantly higher than the background. Comparing with genomic alterations, each of genes connected to more drugs and the top genes had higher cancerous percentages. The top genes with their expression alterations associated with subtypes were different from those with genomic alterations associated with subtypes (Fig. 3D, Fig. 2D). Among them, IKZF1 and IL21R ranked the top two with connections to 164 drugs and 155 drugs (Fig. 3D).

We then investigated the distribution of drugs on their connected genes. The number of genes whose expression was associated with the subtypes varied from 0 to 700 across the drugs, with 14.53% drugs had more than 200 gene connections (Fig. 3E). Moreover, genes whose expression was positively connected with drug resistance across pharmacological subtypes were significantly more abundant than genes that were negatively connected (Fig. 3F). The top 20 drugs with the most connections to genes included UNC1215, affecting the chromatin, NPK76, affecting cell cycle, Methotrexate, affecting DNA replication, AKT inhibitor VIII, affecting PI3K/MTOR signaling pathway, YM201636, affecting PI3K/MTOR signaling pathway, NPK76-II-72-1, affecting the cell cycle pathway, and some compounds such as Zibotentan, A-484954 et al, affecting multiple or unknown pathways (Fig. 3G).

Methotrexate, as an example, is a chemotherapy that specifically acts during DNA and RNA synthesis, and cancer cells were classified into seven subtypes (Fig. 3H) in terms of drug sensitivity to methotrexate. The first subtype, C1, was most sensitive to the drug, and the seventh subtype, C7, was the most resistant to the drug (Fig. 3H). C1 included 12 histological subtypes, with the top 2 which shown in the bottom of Fig. 3H, ALL and DLBC. C2 and C3 were composed of 18 and 19 histological subtypes, respectively, with SCLC both included in the top 2 (Fig. 3H). A total of 297 cancer genes with high expression in sensitive group (C1, C2 and C3) and low expression in resistant group (C4, C5, C6 and C7), which had positive connections to pharmacological subtypes of methotrexate, were selected (Fig. 3I). Pathway enrichment analysis showed that these genes were enriched in pathways including cell cycle and DNA replication et al. (Fig. 3J), suggesting that the ectopic activation of cell cycle lead to the sensitive response of ALL, DLBC, LAML and SLCL to Methotrexate.

### Redefining the similarity of cells based on pharmacological subtypes

Pharmacological subtypes provide us a new point of view to redefine the similarity of cells purely from therapeutic concordance. Thus we calculated the similarity of pairwise cells based on pharmacological subtypes of drugs. If there are quite a few cases that two cells are in the same sensitive or resistant pharmacological subtypes of drugs, these cells gain a high similarity. Otherwise, they gain a low similarity. The similarity of cells defined a hierarchical cluster as shown in Fig. 4A. We further applied the Fisher’s exact test to observe if the cells in the same clusters were from the same histological subtypes (Fig. 4B-D). The points with black border in the figure represent statistical significance (p < 0.05), with the smaller p values shown in red.

**Figure 4.**
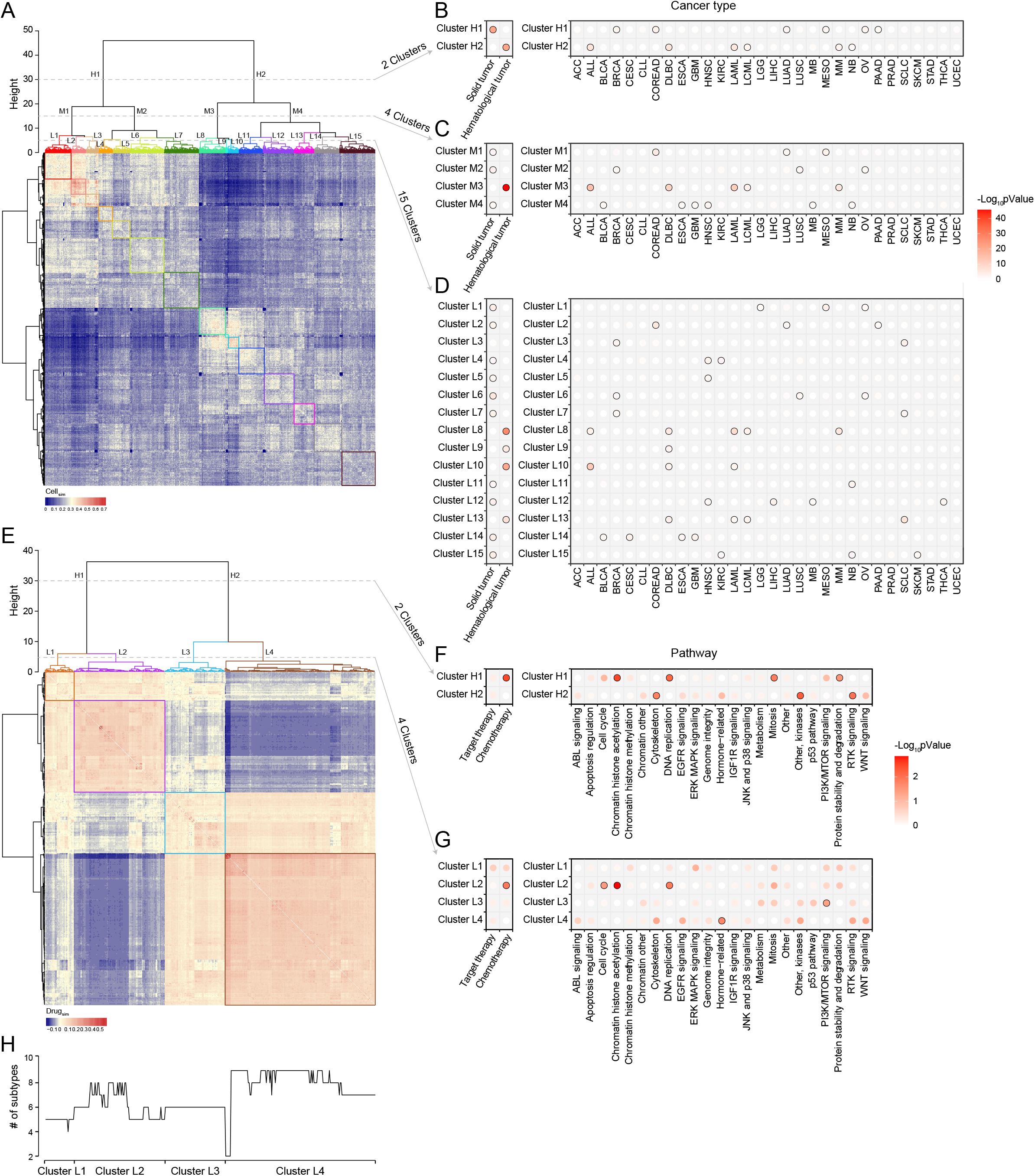
Cell similarity and drug similarity based on pharmacological subtypes. (A) Hierarchical clustering of a cell similarity matrix which was generated based on drug response of cells to drugs, with the red colour indicating high similarity, and the blue colour indicates low similarity between two cell lines. Clusters were formed by partitioning of the dendrogram at a high, median and low height respectively. (B) Two clusters (H1, H2) were formed by cutting the dendrogram at a height of 30. Points with black border represents statistical significance (p < 0.05) determined by Fisher’s exact test. Almost all histological subtypes of blood cancer were grouped together. (C) Four clusters (M1, M2, M3 and M4) were formed by cutting the dendrogram at a height of 15. (D) Fifteen clusters (L1-15) were formed by cutting the dendrogram at a height of 5. Cancers with the same tissue of origin and organ did not cluster together. (E) Hierarchical clustering of the drug similarity profiles with the red colour indicating a high similarity and the blue colour indicating a low level. Clusters were formed by partitioning of the dendrogram at a high and low height respectively. (F) Two clusters (H1, H2) were formed by cutting the dendrogram at a height of 30. Points with black border represents statistical significance (p < 0.05) determined by Fisher’s exact test. Targeted therapy related drugs were grouped together. (G) Four clusters (L1, L2, L3 and L4) were formed by cutting the dendrogram at a height of 5. Cytoskeleton, EGFR signaling, Hormone-related and WNT signalling pathways cluster together. (H) The line chart illustrates the number of subtypes of each drug.

The results showed that the cancer cell lines were respectively divided into 2, 4 and 15 clusters at the different hierarchical levels (Fig. 4A). Two clusters at the high level were reflected by two distinct patterns of similarity (Fig. 4A). One cluster (Cluster H1) was significantly correlated with solid tumors including BRCA, COREAD, LUAD, MESO, OV, and PAAD, and the other (Cluster H2) hematological tumors including ALL, DLBC, LAML, LCML and MM (Fig. 4B). So in a holistic view, the different types of hematological tumors shared a similar drug sensitivity profile, which was quite different from the profile of solid tumors. However, for a given cancer type (the right panel of Fig. 4B), such as ACC, BLCA, CESC, CLL, or ESCA et al., its cells were unbiasedly distributed in two clusters, showing the treatment heterogeneity for the cells from the same tissue origin. At the median hierarchical levels, 4 clusters were identified (Fig. 4C). Among them, Cluster M3 was specifically associated with haematological tumors (p < 0.001, Fisher’s exact test) including ALL, DLBC, LAML, LCML and MM. The treatment heterogeneity was still observed in cancers.

When cell lines were divided into 15 clusters at the lower hierarchical level (Fig. 4D), we observed that the cells from ACC, CLL, STAD, or UCEC were unbiasedly distributed in the different clusters, still showing their treatment heterogeneity. Interestingly, the cells from some cancers such as HNSC, KIRC, and SCLC were enriched in more than 2 clusters with distinctly different drug response. For instance, HNSC cells were enriched in Clusters L12, L4 and L5; KIRC cells were enriched in Clusters L15 and L4; SCLC cells were enriched in Clusters L13, L3 and L7. Interestingly, the SCLC cells in Cluster L13 shared a more similar profile with hematological tumors including DLBC, LAML and LCML. To investigate the underlying mechanism of such treatment heterogeneity in SCLC, we applied DESeq2 analysis to call the differentially expressed genes between Cluster L13 and Cluster L3, and further identified the differential functions via gene set enrichment analysis (GSEA) [16]. The results showed that the functions like DNA replication, RNA transport et al. were significantly upregulated in Cluster L13 cells, while in Cluster L3 cells the functions like immune response, antigen processing and presentation et al. were mainly upregulated (Supple. Fig. 5A). Moreover, MYC, as an oncogenic transcription factor of cell growth and proliferation via enhancing the cell cycle regulated genes [19], was significantly upregulated in Cluster L13 than in L3, with a log2-fold change of 4.28 (Supple. Fig. 5B). These suggested that SCLC cell lines in Cluster L13 were more malignant than those in Cluster L3.

### Redefining the similarity of drugs based on pharmacological subtypes

The similarity of pairwise drugs was also defined as the summarized similarity of drug response across the same cells which were tested for both drugs. For two drugs, if there are quite a few of cell pairs which are in the sensitive (or resistant) pharmacological subtypes of one drug are also in the sensitive (or resistant) subtypes of the other drug, they gain a high similarity. Otherwise, they gain a low similarity. We then generated a drug similarity profile across 367 drugs, and similarly, a hierarchical clustering algorithm was used to group drugs based on drug similarity (Fig. 4E). The results showed that at the high level, drugs were divided into two clusters, demonstrating two different similarity patterns. The first cluster (Cluster H1) mostly included chemotherapy drugs such as those effecting in DNA replication and mitosis, and the second cluster was mixed with target and chemotherapy drugs, specifically enriched by the drugs on RTK signaling, other kinases, and cytoskeleton (Fig. 4F). When the drugs were divided into 4 clusters (Fig. 4E, G), we observed that the drugs enriched in L2 included chemotherapy drugs effecting in cell cycle, chromatin histone accelylation, and DNA replication, the drugs enriched in L3 were functioning in PI3K/mTOR signaling pathway, and the drugs enriched in L4 were hormone related ones. Moreover, L4, accounting for a massive part of drugs, also included the drugs targeting the different types of signaling pathways such as ABL, EGFR, Jak and P38, RTK and WNT signalling. These drugs did not reach the significant level but provided the obvious association.

We further observed the number of pharmacological subtypes of these drugs that were arranged by the clustering structure above (Fig. 4H). Remarkably, there was the varied treatment heterogeneity for the drugs in the different clusters. The drugs in Cluster L4 had more subtypes but those in Cluster L1 and L3 had less subtypes, suggesting higher treatment heterogeneity for L4 and lower treatment heterogeneity for L1 and L3.

### An access to pharmacological subtypes of drugs

We developed a website to store and visualize the pharmacological subtypes of drugs, as well as the distributions of histological subtypes across them, and connected genomic and expression alterations. All data and results used for the study are easily accessible via http://www.hywanglab.cn/dtdb/ (Supple. Fig. 6).

## Discussion

In this study, we integrated drug response information to pharmacologically reclassify tumour subtypes, as well as identify intrinsic concordant molecular mechanisms. Unlike the many studies that have aimed to unravel cancer heterogeneity by defining subtypes within the same tissues, our study aimed to systematically uncover that cancers across disparate tissues of origin can belong to one pharmacological subtype, benefiting from similar anticancer therapies, because they share a common molecular mechanism of oncogenesis. Besides, we also developed the new measures to redefine cell similarity and drug similarity from the therapeutic concordance, which provided a new point of view to study cancer heterogeneity. The similarity of cells further depicted that cells from the different origin of tissue could share the similar responses of drugs; likewise, that cells from the same origin of tissue could have distinct drug responses, thus indicating the new subtypes.

Our analysis identified, for instance, that although SKCM and COREAD are histologically different, both belonged to one pharmacological subtype, the MEK1/2 inhibitor C1 subtype, and were similarly sensitive to MEK1/2 inhibitors (Fig. 1A-D). By connecting genomic alterations with pharmacological subtypes, mutations in the oncogene BRAF or KRAS were found to be overwhelmingly more frequent in the C1 group (Fig. 2H). BRAF and KRAS are the kinases upstream of MEK1/2 that transmits the signals down through MEK1/2 without other major signalling branches [20, 21]. Our results illustrated that MEK inhibitors were able to effectively block aberrantly activated signals from BRAF or KRAS mutations. Further investigation by cBioportal (https://www.cbioportal.org/) of TCGA patients also revealed that approximately 60% of THCA (thyroid carcinomas) and 50% of SKCM (melanomas) harboured BRAF mutations, and 65% of PAAD (pancreas) and 40% COREAD (colorectal carcinomas) possessed KRAS mutations, a rate that was strikingly higher than that of other cancers (Supple. Fig. 7A, B). Given the high mutant frequency of BRAF in THCA and SKCM, and KRAS in PAAD and COREAD, such 4 cancer types were significantly enriched in C1 pharmacological subtype of MEK1/2 inhibitor PD0325901 (THCA, *p*=8.79e-4; SKCM, *p*=1.30e-14; PAAD, *p*=4.17e-3; COREAD, *p*=4.58e-4; Hypergeometric Distribution Test) (Supple. Tab. 2). In contrast, mutations in the oncogenes RB1 was found to be less frequent in the C1 group than in other groups. RB1 is a negative regulator of the cell cycle, with its active hypophosphorylated form binding the transcription factor E2F1 [22]. RB1 mutations lead to the ectopic activation of the cell cycle, which cannot be controlled using MEK inhibitors. This may be because RB1, located downstream of MEK1/2, gain ectopic activation independent of upstream stimulation. In another instance, therapeutic concordance was identified across histologically disparate blood cancers in regard to methotrexate, a chemotherapy drug specially acting during DNA and RNA synthesis (Fig. 3H). This is well known that Methotrexate is used in haematological malignancies but less so in solid cancers [23].

Functional analysis of the positively correlated genes showed that haematological tumors had more active gene expression related to the cell cycle than solid tumours (Fig. 3I, J). The molecular basis of such extraordinary activation of the cell cycle in haematological tumors could explain their good response to chemotherapy drugs, which work mainly by inhibiting mitosis and cell division. This demonstrates pharmacological subtypes benefiting from similar anticancer therapies can be due to common molecular mechanisms related to tissue-specific gene expression and pathways. Our website provided the pharmacological subtypes of 367 drugs as well as their connections with the molecular alterations for the users to retrieve and download.

We also investigated the treatment heterogeneity for six types of histological cancers, including LUAD, BRCA, SKCM, SCLC, COREAD and HNSC. High treatment heterogeneity in LUAD was observed compared with other types of cancers, suggesting that anticancer therapies for LUAD should be more complicated than those for other types of cancers (Fig. 1F-H). Moreover, the increased number of subtypes of EGFR inhibitors in LUAD indicated there was high treatment heterogeneity when EGFR inhibitors were applied into the therapies of LUAD patients (Fig. 1H).

Our analysis showed that genes whose genomic/expression alterations frequently connect to anticancer drugs had a higher likelihood of being cancer genes. At genomic level, the top 10 genes were connected to more than 9 drugs, 20% of which (including BRAF and KRAS) were annotated as cancer censor genes. At expression level, each of genes connected to more drugs and the top genes had higher cancerous percentages. This indicated that some genes currently not identified as cancer genes could actually be causally implicated in cancer. In our analysis, ADK-VCL, TNFRSF9 and LTB, as the top three most connected genes, were identified as novel cancer genes for further investigation. Adenosine kinase (ADK) has an important role in mitosis, tumorigenesis, tumor-associated tissue remodeling, and invasion, and manipulation of ADK has the potential to be a therapeutic strategy for invasive breast cancer [24]. Vinculin (VCL) is a cytoskeletal protein associated with cell-cell and cell-matrix junctions, which interacts with many focal adhesion proteins and has been found to be closely linked to tumor migration [25]. ADK-VCL had 45 drug connections suggesting its therapeutic potential, although ADK-VCL fusion events were rarely reported. We speculated that its fusion was beneficial for tumor metastasis, but evidence was lacking. The tumor necrosis factor receptor superfamily member 9 (TNFRSF9) with connections to 25 drugs, also known as 4-1BB and CD137, is an immune co-stimulatory receptor. TNFRSF9 is expressed on activated immune cells including natural killer (NK) cells, effector T cells and antigen presenting cells, among them dendritic cells, macrophages, and B cells [26]. LTB (Lymphotoxin Beta), with connections to 20 drugs, is a type II membrane protein of the TNF family. Giuseppina et al. found that a decrease in lymphotoxin-β production by tumor cells was associated with a loss of follicular dendritic cell phenotype and diffuse growth of follicular lymphomas [27]. Recently, the gene was found to be associated with immune infiltration of breast and endometrial cancer tumors [28, 29]. The analysis using the TCGA dataset showed that TNFRSF9 was altered in approximately 5% of cases of Cholangiocarcinoma, adrenocortical carcinoma (ACC) and lymphoid neoplasm diffuse large B-cell lymphoma (DLBC), with deep deletion being the dominant alteration (Supple. Fig. 8A). In addition, more than 14% of DLBC patients had altered LTB genes, with mutations and deletion being the major factors (Supple. Fig. 8B). Therefore, the importance of the roles of these genes in cancers could be underappreciated.

Genes whose alterations were positively connected with drug resistance were significantly more frequent than those negatively connected at the mRNA expression level but not at the genomic level (Fig. 3F). A positive connection meant that genomic alterations or high expression indicated increased drug sensitivity. In this way, genes with a positive connection could be indicators of therapeutic efficiency for connected drugs. Genes with negative connections could be potential biomarkers for developing new therapies to overcome the resistance of connected drugs. Our analysis suggests that most of the genes whose expressions are altered during cancer initiation and development (driver genes or not) are located in ectopically activated pathways that could be controlled by anticancer drugs. Only a proportion of altered genes are outside of the cellular pathways targeted by anticancer drugs, and their alterations may maintain the growth signals of cancer cells when anticancer drugs are used. Therefore, these genes may be potential therapeutic biomarkers for overcoming drug resistance to the connected drugs.

Based on pharmacological subtypes, the cell similarity and the drug similarity could be re-defined (Fig. 4). The cells from the same origin of tissue can be dispersed in the different pharmacological subtypes, showing the completely different response to the same drug, because of their intrinsically molecular heterogeneity. For instance, HNSC, KIRC and SLCL cells were separated into more than two clusters with distinct drug response (Fig. 4D). Specifically, the SCLC cells with more malignant signatures, in particular, higher expression levels of MYC and upregulated pathways involving DNA replication and cell cycle, were far away from the other SCLC cells and close to hematological tumors, instead. Interestingly, by connecting to expression alterations, our analysis further confirmed that SCLC cells more like hematological tumors had the characteristic of the ectopic activation of cell cycle, which led to their sensitive to a chemotherapy Methotrexate (Fig. 3H-J). Our pharmacological analysis has recognized a new subtype of SCLC cells, with the high expression of an oncogenic transcription factor MYC as a marker, can benefit from chemotherapy than other SCLC subtypes. This finding was consistent with the latest studies [30, 31]. Therefore, the subtypes identified by re-classifying the same origin of tissue using pharmacological data are worth to be further investigated. In addition, our analysis of the drug similarity unraveled that the drugs were categorized into two major groups, wherein the chemotherapeutic drugs tended to share the similar pattern (Fig. 4E, F). Moreover, Cluster L4 including many drugs targeting the signalling pathways had more pharmacological subtypes, suggesting a high treatment heterogeneity for some target drugs (Fig. 4H).

## Conclusions

In summary, our study was a systematic analysis which aimed to reveal intrinsic therapeutic concordance across histologically disparate tumour subtypes. The findings provide us with a purely therapy-oriented perspective to re-examine tumour classifications independent of histology subtypes.

## Availability of data and materials

The data and materials that supporting the conclusion of this paper have been included within the article and the website (www.hywanglab.cn/dtdb/).

## Acknowledgements

This work was supported by grants from the National Key Research and Development Program (2017YFC0908500 to HW) and the National Natural Science Foundation of China (31771469 and 31571363 to HW; 81772465 to FZ).

## Authors’ contributions

HW conceived the hypothesis. YX, JZ and HW designed and performed the data analysis. ZC, WL, YD, XC and TW collected and preprocessed the data. HW, YX, JZ, JK and FZ wrote and revised the manuscript.

## Supplementary Data

**Supplementary Figure 1. The comparisons between two types of cancers by the numbers of pharmacological subtypes across drugs.**

**Supplementary Figure 2. The top 30 genes whose genomic alterations frequently connect to pharmacological subtypes.**

**Supplementary Figure 3. Genomic alterations connect to pharmacological subtypes based on MEK inhibitors, Refametinib, Trametinib, Selumetinib, PD-0325901, and CI-1040.** BRAF mutation and KRAS mutation were positively connected to 5 and 4 drugs, respectively. RB1 mutation had a negative connection to 5 drugs.

**Supplementary Figure 4. The top 30 genes whose expressions frequently connect to pharmacological subtypes.**

**Supplementary Figure 5. Pathways significantly enriched in the comparison of Cluster L13 vs. L3 and the different levels of MYC expression in such comparison.**

**Supplementary Figure 6. Introduction of the website.**

**Supplementary Figure 7. Mutation frequency of BRAF and KRAS in pan-cancer patients in the TCGA dataset.** (A) 60% of thyroid carcinomas and 50% of melanomas harboured BRAF mutation. (B) 65% of pancreas and 40% colorectal carcinomas harboured KRAS mutation.

**Supplementary Figure 8. Mutation frequency of TNFRSF9 and LTB in pan-cancer patients in the TCGA dataset.** (A) TNFRSF9 was altered in cholangiocarcinoma, lymphoid neoplasm diffuse large B-cell lymphoma, adrenocortical carcinoma, etc. in approximately 5% of cases, with deep deletion being the dominant alteration. (B) More than 14% of DLBC patients had altered LTB genes.

**Supplementary Table 1. Genes mapped to core cancer pathways.**

**Supplementary Table 2. The enrichment of histological cancers in the most sensitive pharmacological subtype of PD0325901.**

## Abbreviations

ACC: Adrenocortical carcinoma
ALL: Acute lymphoblastic leukemia
BLCA: Bladder urothelial carcinoma
BRCA: Breast invasive carcinoma
CESC: Cervical squamous cell carcinoma and endocervical adenocarcinoma
CLL: Chronic lymphocytic leukemia
COREAD: Colon adenocarcinoma and rectum adenocarcinoma
DLBC: Lymphoid neoplasm diffuse large B-cell Lymphoma
ESCA: Esophageal carcinoma
GBM: Glioblastoma multiforme
HNSC: Head and neck squamous cell carcinoma
KIRC: Kidney renal clear cell carcinoma
LAML: Acute myeloid leukemia
LCML: Chronic myelogenous leukemia
LGG: Brain lower grade glioma
LIHC: Liver hepatocellular carcinoma
LUAD: Lung adenocarcinoma
LUSC: Lung squamous cell carcinoma
MB: Medulloblastoma
MESO: Mesothelioma
MM: Multiple myeloma
NB: Neuroblastoma
OV: Ovarian serous cystadenocarcinoma
PAAD: Pancreatic adenocarcinoma
PRAD: Prostate adenocarcinoma
SCLC: Small cell lung cancer
SKCM: Skin cutaneous melanoma
STAD: Stomach adenocarcinoma
THCA: Thyroid carcinoma
UCEC: Uterine corpus endometrial carcinoma
CGP: The Cancer Genome Project
IC50: Half maximal inhibitory concentration
TCGA: The Cancer Genome Atlas Program
FDA: U.S. Food and Drug Administration
RTK: Receptor tyrosine kinase
MEK: Mitogen-activated protein kinase kinase

## References

1. Swanton, C., Intratumor heterogeneity: evolution through space and time. Cancer Res, 2012. 72(19): p. 4875–82.

2. Bedard, P.L., et al., Tumour heterogeneity in the clinic. Nature, 2013. 501(7467): p. 355–64.

3. Vogelstein, B., et al., Cancer genome landscapes. Science, 2013. 339(6127): p. 1546–58.

4. Tran, B., et al., Cancer genomics: technology, discovery, and translation. J Clin Oncol, 2012. 30(6): p. 647–60.

5. Chapman, P.B., et al., Improved survival with vemurafenib in melanoma with BRAF V600E mutation. N Engl J Med, 2011. 364(26): p. 2507–16.

6. Mok, T.S., et al., Gefitinib or carboplatin-paclitaxel in pulmonary adenocarcinoma. N Engl J Med, 2009. 361(10): p. 947–57.

7. Perou, C.M., et al., Molecular portraits of human breast tumours. Nature, 2000. 406(6797): p. 747–52.

8. Cancer Genome Atlas, N., Comprehensive molecular portraits of human breast tumours. Nature, 2012. 490(7418): p. 61–70.

9. Prat, A., et al., PAM50 assay and the three-gene model for identifying the major and clinically relevant molecular subtypes of breast cancer. Breast Cancer Res Treat, 2012. 135(1): p. 301–6.

10. Schmitz, R., et al., Genetics and Pathogenesis of Diffuse Large B-Cell Lymphoma. N Engl J Med, 2018. 378(15): p. 1396–1407.

11. Cancer Genome Atlas Research, N., et al., The Cancer Genome Atlas Pan-Cancer analysis project. Nat Genet, 2013. 45(10): p. 1113–20.

12. Hoadley, K.A., et al., Multiplatform analysis of 12 cancer types reveals molecular classification within and across tissues of origin. Cell, 2014. 158(4): p. 929–944.

13. Drilon, A., et al., Efficacy of Larotrectinib in TRK Fusion-Positive Cancers in Adults and Children. N Engl J Med, 2018. 378(8): p. 731–739.

14. Garnett, M.J., et al., Systematic identification of genomic markers of drug sensitivity in cancer cells. Nature, 2012. 483(7391): p. 570–5.

15. Benjamini, Y. and Y. Hochberg, Controlling the False Discovery Rate: A Practical and Powerful Approach to Multiple Testing. Journal of the Royal Statistical Society: Series B (Methodological), 1995. 57(1): p. 289–300.

16. Subramanian, A., et al., Gene set enrichment analysis: a knowledge-based approach for interpreting genome-wide expression profiles. Proc Natl Acad Sci U S A, 2005. 102(43): p. 15545–50.

17. Love, M.I., W. Huber, and S. Anders, Moderated estimation of fold change and dispersion for RNA-seq data with DESeq2. Genome Biol, 2014. 15(12): p. 550.

18. Tate, J.G., et al., COSMIC: the Catalogue Of Somatic Mutations In Cancer. Nucleic Acids Res, 2019. 47(D1): p. D941–d947.

19. Oster, S.K., et al., The myc oncogene: MarvelouslY Complex. Adv Cancer Res, 2002. 84: p. 81–154.

20. McCubrey, J.A., et al., Roles of the Raf/MEK/ERK pathway in cell growth, malignant transformation and drug resistance. Biochim Biophys Acta, 2007. 1773(8): p. 1263–84.

21. Simanshu, D.K., D.V. Nissley, and F. McCormick, RAS Proteins and Their Regulators in Human Disease. Cell, 2017. 170(1): p. 17–33.

22. Chen, H.Z., S.Y. Tsai, and G. Leone, Emerging roles of E2Fs in cancer: an exit from cell cycle control. Nat Rev Cancer, 2009. 9(11): p. 785–97.

23. Kożmiński, P., et al., Overview of Dual-Acting Drug Methotrexate in Different Neurological Diseases, Autoimmune Pathologies and Cancers. Int J Mol Sci, 2020. 21(10).

24. Shamloo, B., et al., Dysregulation of adenosine kinase isoforms in breast cancer. Oncotarget, 2019. 10(68): p. 7238–7250.

25. de Semir, D., et al., PHIP drives glioblastoma motility and invasion by regulating the focal adhesion complex. Proc Natl Acad Sci U S A, 2020. 117(16): p. 9064–9073.

26. Fröhlich, A., et al., Comprehensive analysis of tumor necrosis factor receptor TNFRSF9 (4-1BB) DNA methylation with regard to molecular and clinicopathological features, immune infiltrates, and response prediction to immunotherapy in melanoma. EBioMedicine, 2020. 52: p. 102647.

27. Pepe, G., et al., Reduced lymphotoxin-beta production by tumour cells is associated with loss of follicular dendritic cell phenotype and diffuse growth in follicular lymphoma. J Pathol Clin Res, 2018. 4(2): p. 124–134.

28. Terkelsen, T., et al., Secreted breast tumor interstitial fluid microRNAs and their target genes are associated with triple-negative breast cancer, tumor grade, and immune infiltration. Breast Cancer Res Treat, 2020. 22(1): p. 73.

29. Ding, H., et al., Prognostic Implications of Immune-Related Genes’ (IRGs) Signature Models in Cervical Cancer and Endometrial Cancer. Front Genet, 2020. 11: p. 725.

30. Bian, X., et al., The MYC Paralog-PARP1 Axis as a Potential Therapeutic Target in MYC Paralog-Activated Small Cell Lung Cancer. Front Oncol, 2020. 10: p. 565820.

31. Ireland, A.S., et al., MYC Drives Temporal Evolution of Small Cell Lung Cancer Subtypes by Reprogramming Neuroendocrine Fate. Cancer Cell, 2020. 38(1): p. 60–78.e12.

