## Supplementary figures and images for "Drug-driven reclassification of multiple tumour subtypes reveals intrinsic molecular concordance of therapy across histologically disparate cancers"

### Supple Figure 1

#### Supple Figure 1

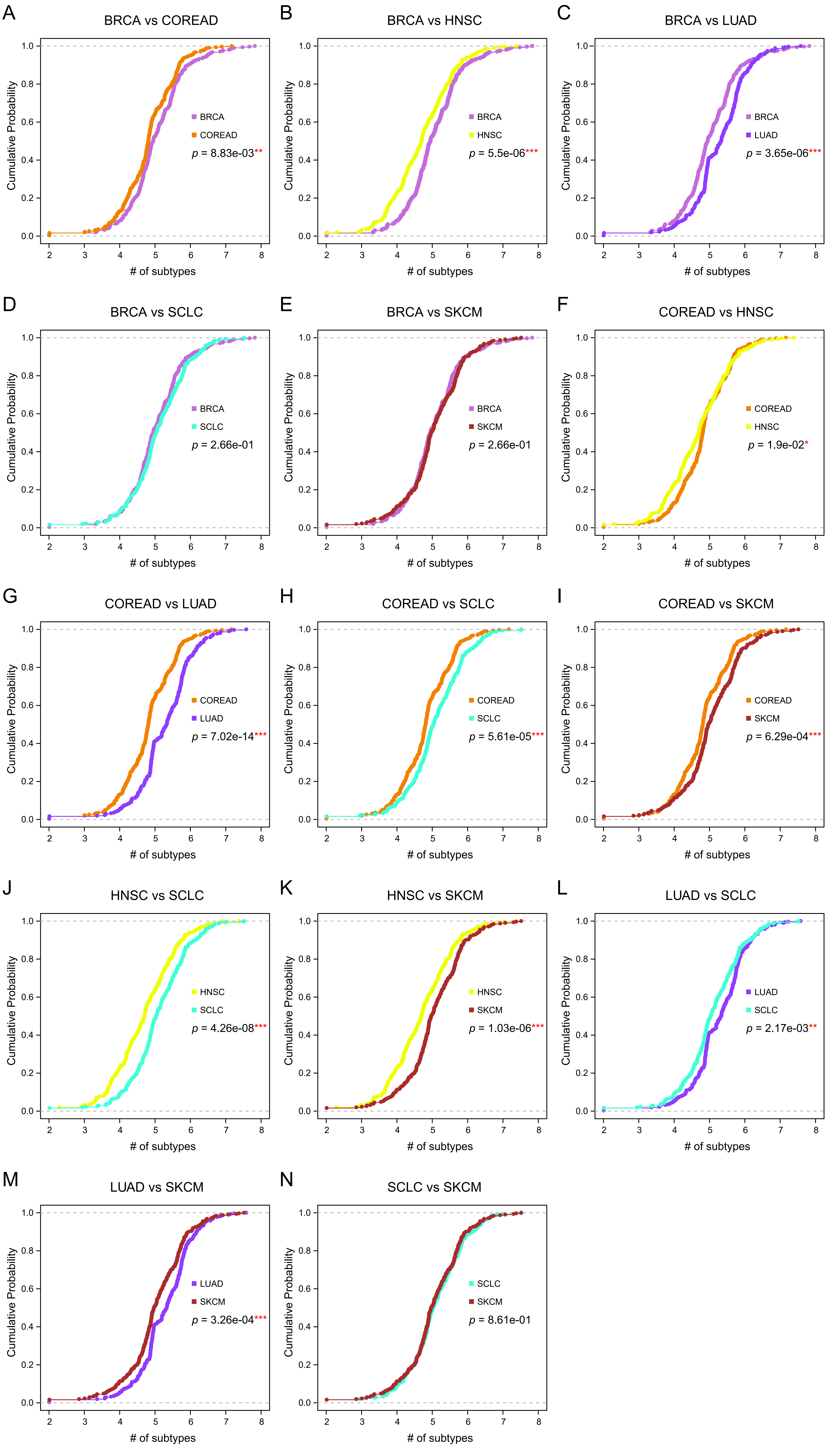

### Supple Figure 3

#### Supple Figure 3

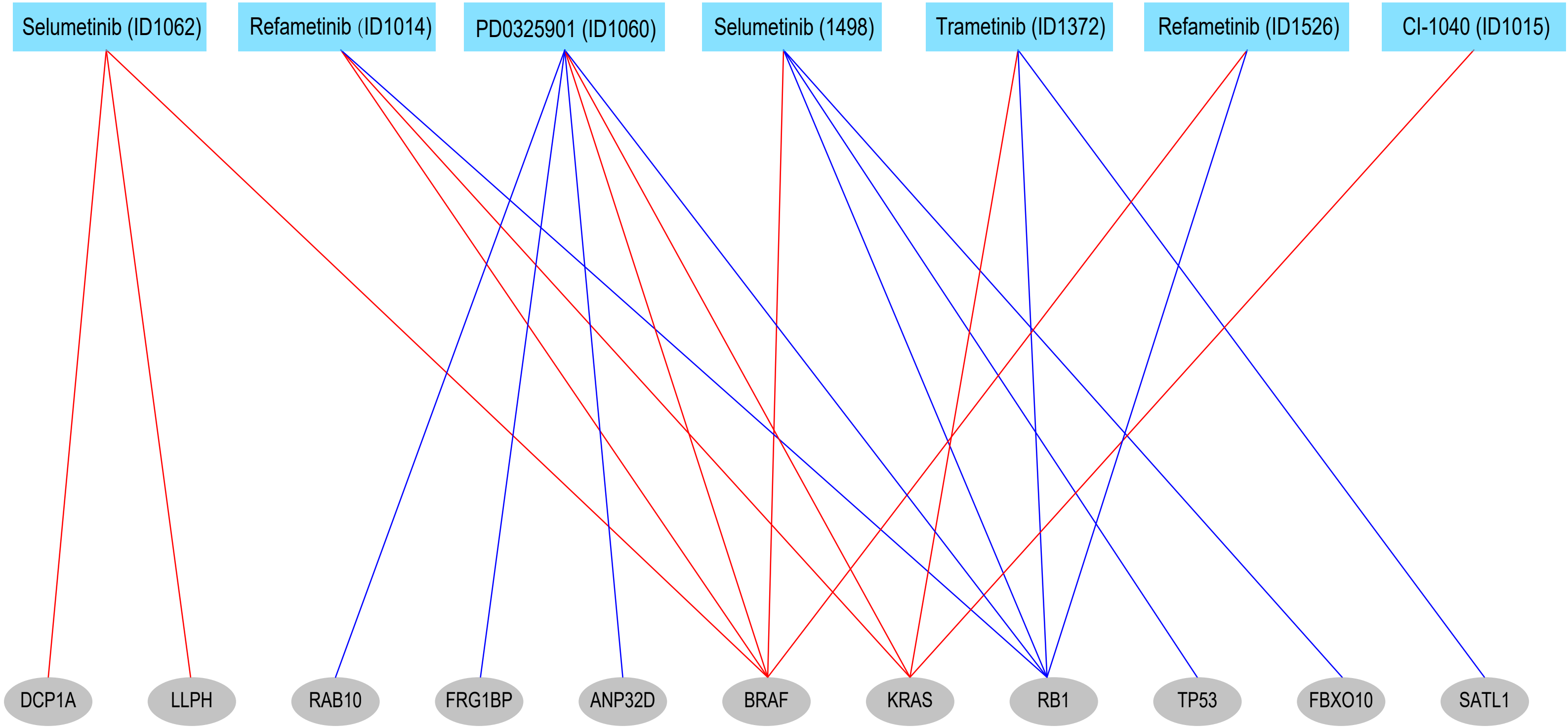

### Supple Figure 5

#### Supple Figure 5

A

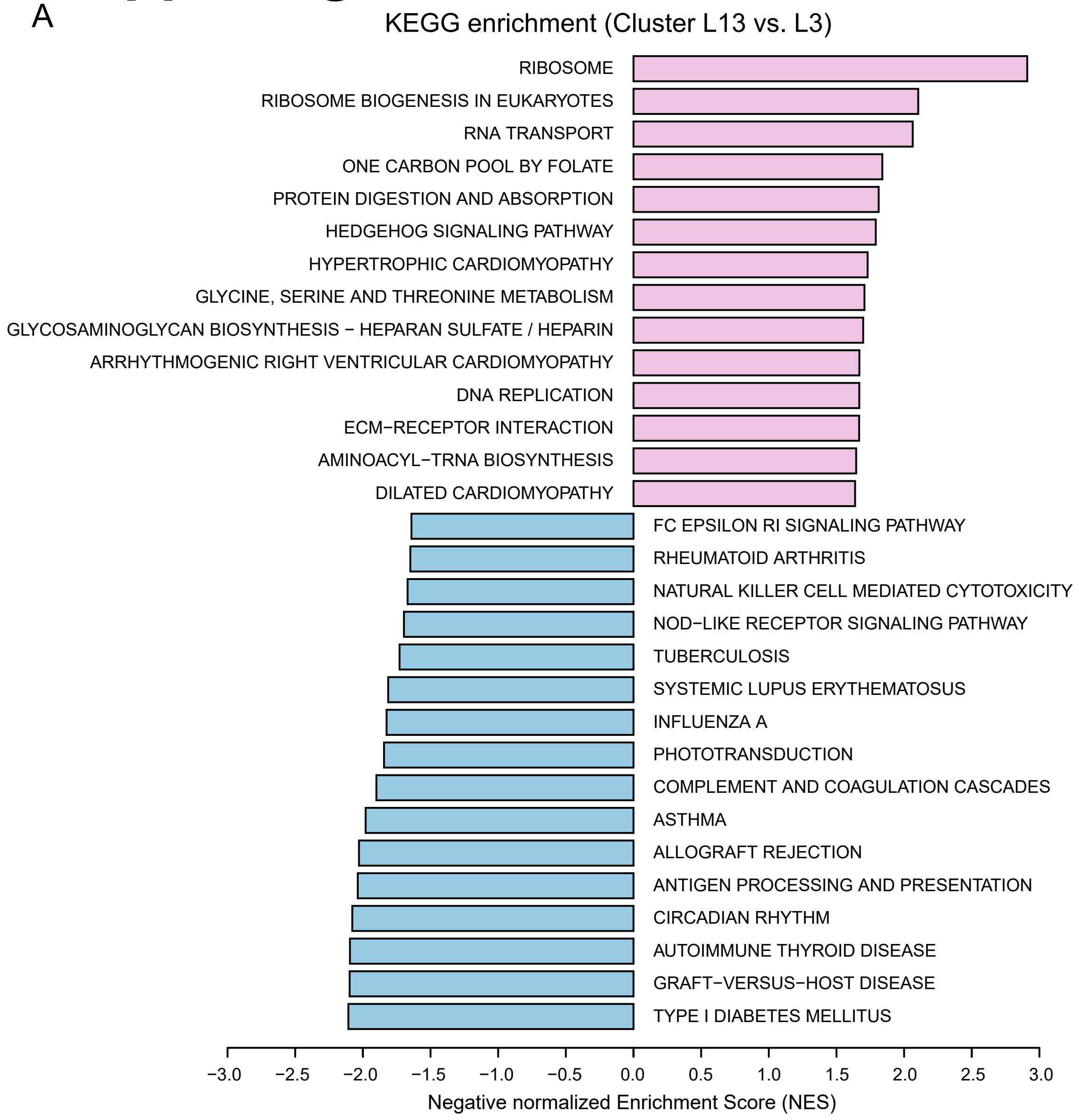


B

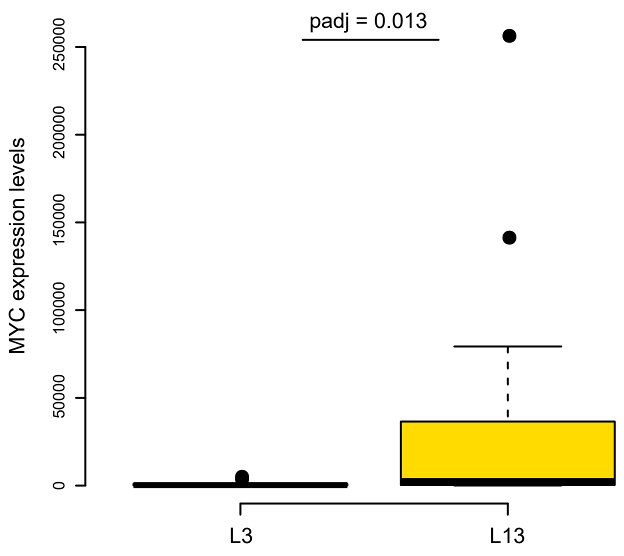

### Supple Figure 6

#### Supple Figure 6

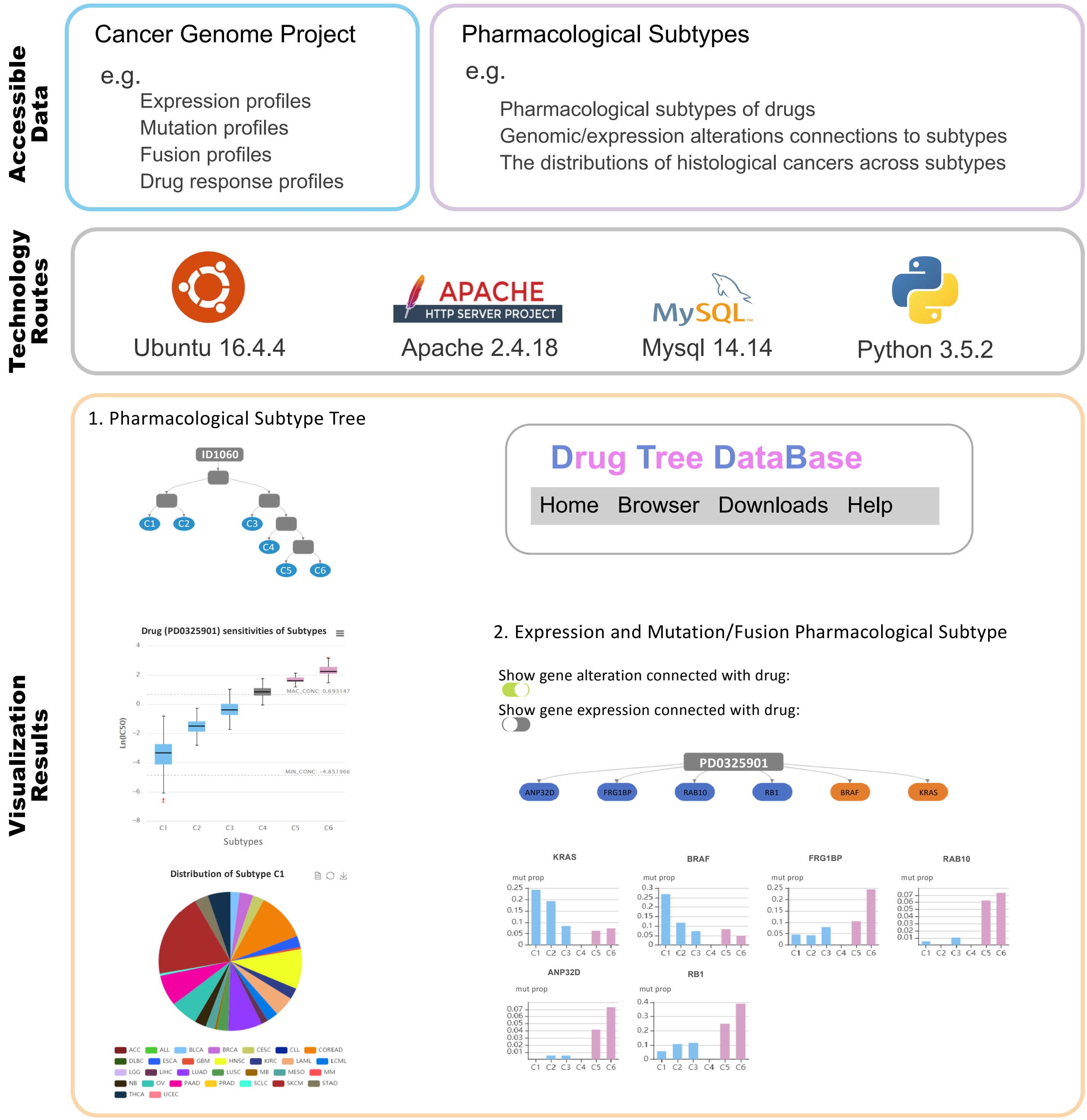

### Supple Figure 7

#### Supple Figure 7

A

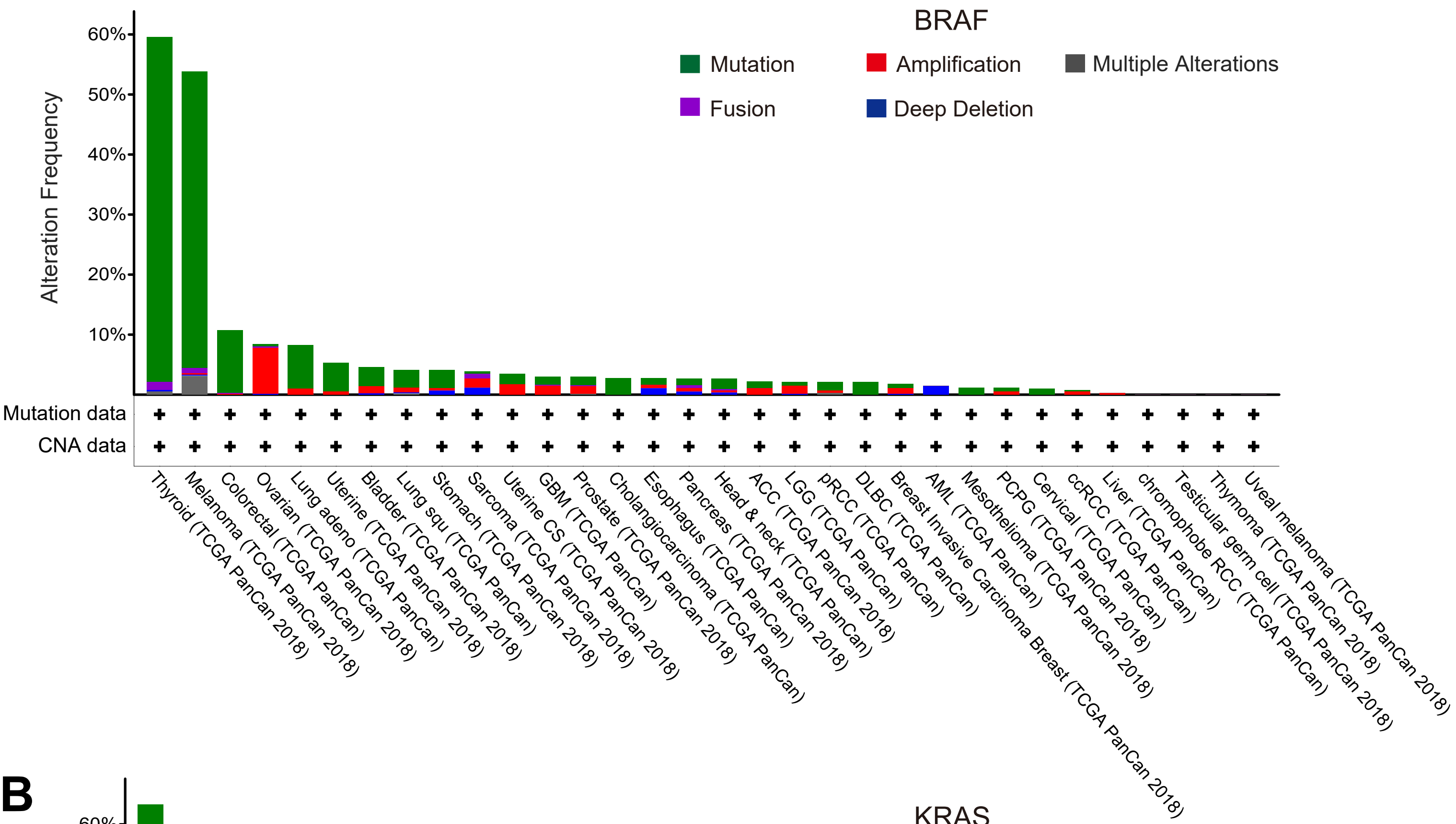


B

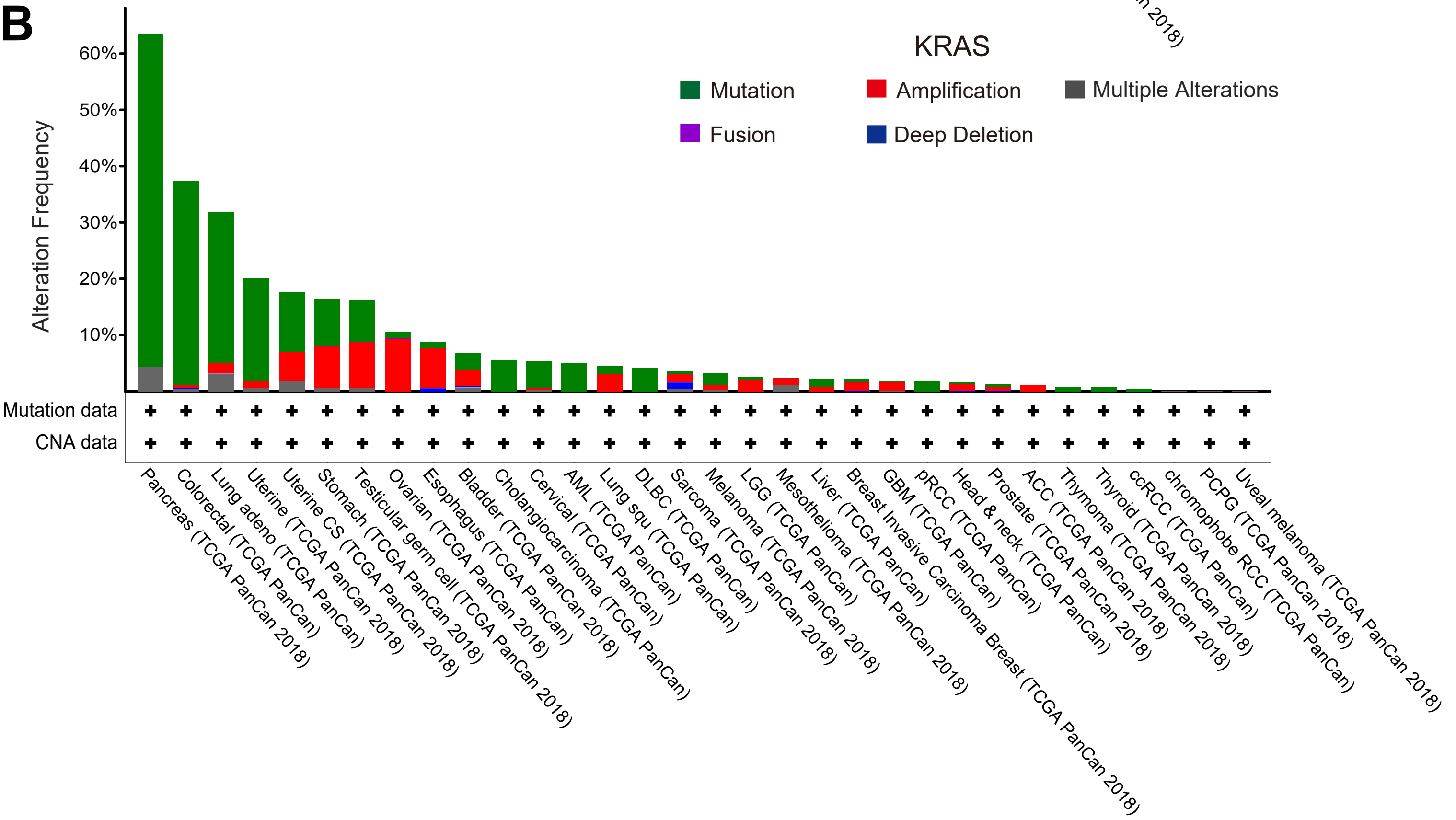

### Supple Figure 8

#### Supple Figure 8

A

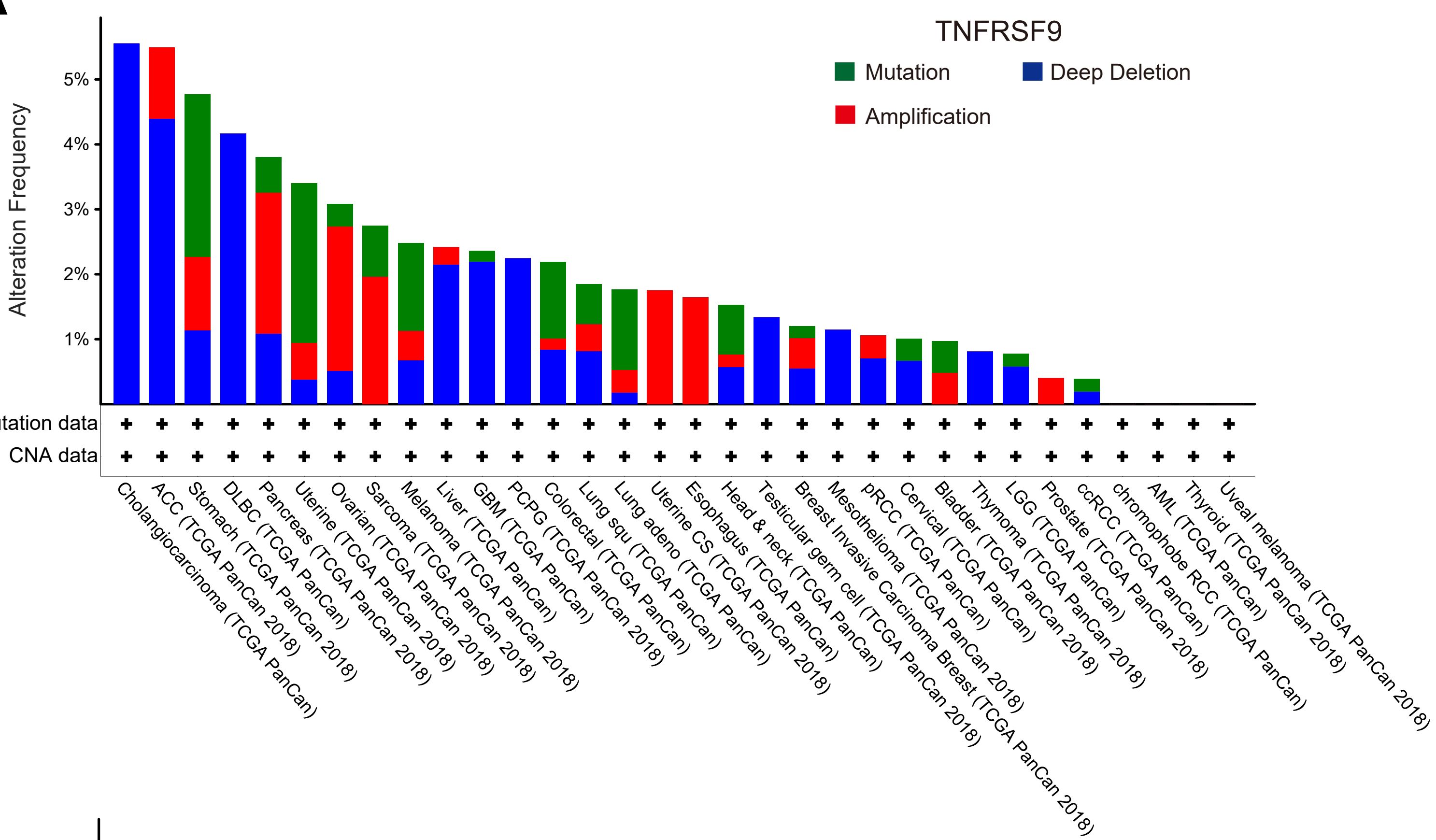


B

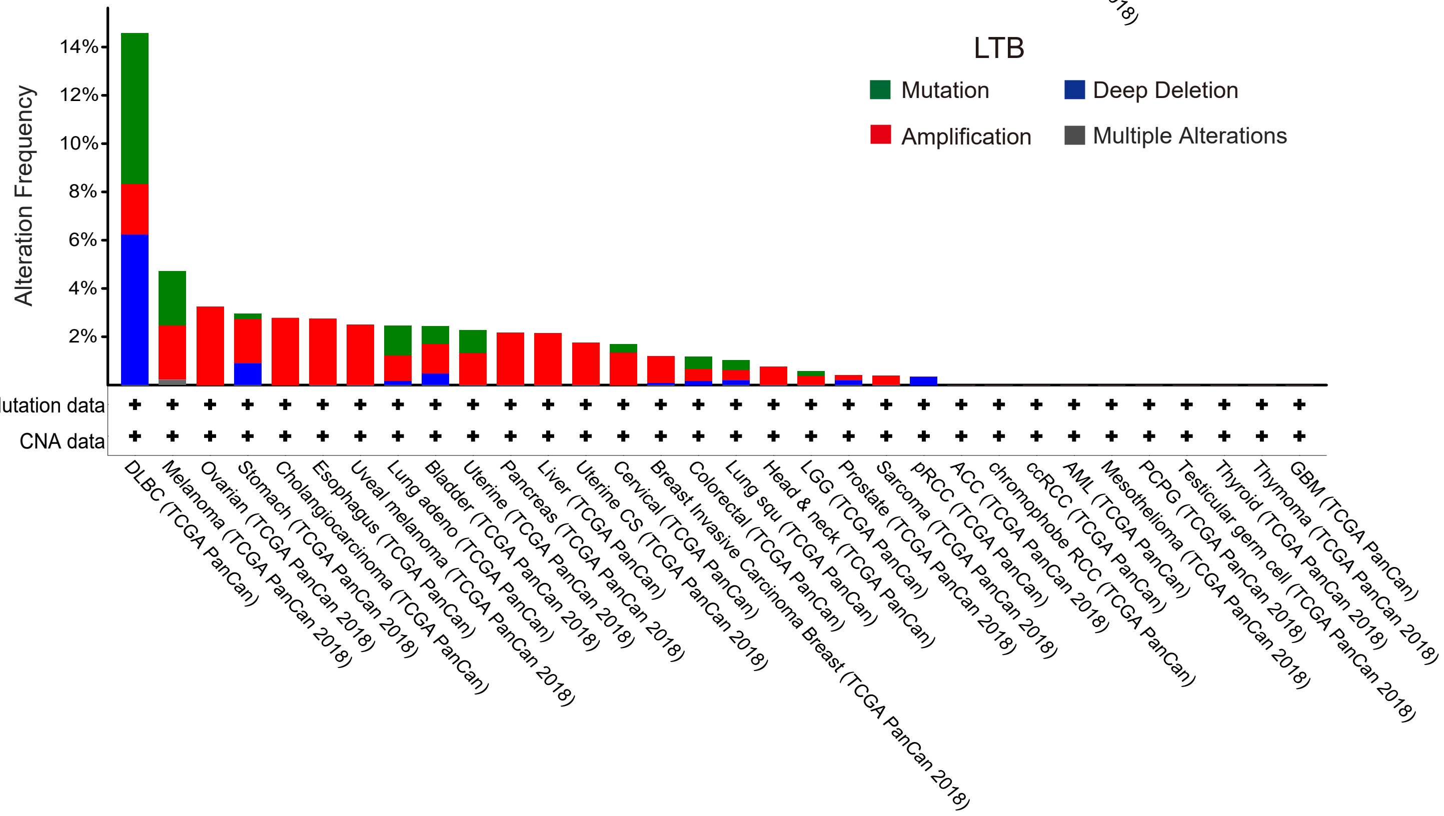
