## Supplementary material for "Drug-driven reclassification of multiple tumour subtypes reveals intrinsic molecular concordance of therapy across histologically disparate cancers": Supple Figure 2

| Gene | # of connection | Cancer gene |
|---|---|---|
| ADK-VCL | 45 | No |
| SEMA3F-CYB561D2 | 27 | No |
| LYZL6-KIAA0930 | 26 | No |
| TNFRSF9 | 25 | No |
| LTB | 20 | No |
| KPRP | 16 | No |
| BRAF | 13 | Yes |
| NFATC4 | 12 | No |
| KRAS | 11 | Yes |
| NREP-AS1-WDR36 | 9 | No |
| TP53 | 9 | Yes |
| HLA-B | 9 | No |
| SEL1L-STON2 | 8 | No |
| BTG2 | 8 | No |
| ZBTB7A | 7 | No |
| SYNGR1 | 7 | No |
| TMSB4XP1 | 7 | No |
| RB1 | 7 | Yes |
| MAFG | 7 | No |
| FOXC2 | 6 | No |
| EMD | 6 | No |
| MRPS18A | 6 | No |
| PTMA | 6 | No |
| FAM229B | 6 | No |
| TMEM121 | 6 | No |
| GRAP2 | 6 | No |
| SLC15A3 | 6 | No |
| DERA | 6 | No |
| SUGP1 | 6 | No |
| SLC39A7 | 6 | No |
