## Supplementary material for "Drug-driven reclassification of multiple tumour subtypes reveals intrinsic molecular concordance of therapy across histologically disparate cancers": Supple Figure 4

| Gene | # of connection | Cancer gene |
|---|---|---|
| PIK3CG | 168 | No |
| IL12RB1 | 167 | No |
| OSM | 167 | No |
| PTPN7 | 167 | No |
| IKZF1 | 164 | Yes |
| IL21R | 155 | Yes |
| RASGRP2 | 155 | No |
| TRAT1 | 150 | No |
| GNGT2 | 145 | No |
| PRKCB | 143 | Yes |
| IL9R | 142 | No |
| CSF2RB | 141 | No |
| CD19 | 140 | No |
| CD28 | 138 | Yes |
| MAP4K1 | 138 | No |
| TNF | 134 | No |
| PIK3R5 | 133 | No |
| IL2RB | 132 | No |
| CD86 | 131 | No |
| FCER2 | 129 | No |
| IL10RA | 129 | No |
| PLCB2 | 126 | No |
| IL2RG | 124 | No |
| PSMA8 | 118 | No |
| BTC | 117 | No |
| JAK3 | 109 | Yes |
| ESR2 | 106 | No |
| INHBB | 104 | No |
| VAV1 | 103 | Yes |
| IL20RA | 102 | No |
