## Supplementary material for "Drug-driven reclassification of multiple tumour subtypes reveals intrinsic molecular concordance of therapy across histologically disparate cancers": Supple Table 1

Supplementary Table 1. Genes mapped in core cancer pathways annotated by KEGG (Kyoto Encyclopedia of Genes and Genomes), MIPS (The Munich Information Center for Protein Sequences), BIOCARTA, PID (Pathway Interaction Database), and REACTOME databases.

| | |
|---|---|
| BIOCARTA TGFB PATHWAY | EP300,MAP2K1,APC,MAP3K7,ZFYVE9,TGFB2,TGFB1,CREBBP,MAPK3,TAB1,SMAD3,SMAD4,TGFBR2,SKIL,TGFBR1,SMAD7,TGFB3,CDH1,SMAD2 |
| KEGG TGF BETA SIGNALING PATHWAY | TFDP1,NOG,TNF,GDF7,INHBB,INHBC,COMP,INHBA,THBS4,RHOA,CREBBP,ROCK1,ID1,ID2,RPS6KB1,RPS6KB2,CUL1,LOC728622,ID4,SMAD3,MAPK3,RBL2,SMAD4,RBL1,NODAL,SMAD1,MYC,SMAD2,MAPK1,SMURF2,SMURF1,EP300,BMP8A,GDF5,SKP1,CHRD,TGFB2,TGFB1,IFNG,CDKN2B,PPP2CB,PPP2CA,PPP2R1A,ID3,SMAD5,RBX1,FST,PITX2,PPP2R1B,TGFBR2,AMHR2,LTBP1,LEFTY1,AMH,TGFBR1,SMAD9,LEFTY2,SMAD7,ROCK2,TGFB3,SMAD6,BMPR2,GDF6,BMPR1A,BMPR1B,ACVRL1,ACVR2B,ACVR2A,ACVR1,BMP4,E2F5,BMP2,ACVR1C,E2F4,SP1,BMP7,BMP8B,ZFYVE9,BMP5,BMP6,ZFYVE16,THBS3,INHBE,THBS2,DCN,THBS1, |
| KEGG WNT SIGNALING PATHWAY | JUN,LRP5,LRP6,PPP3R2,SFRP2,SFRP1,PPP3CC,VANGL1,PPP3R1,FZD1,FZD4,APC2,FZD6,FZD7,SENP2,FZD8,LEF1,CREBBP,FZD9,PRICKLE1,CTBP2,ROCK1,CTBP1,WNT9B,WNT9A,CTNNBIP1,DAAM2,TBL1XR1,MMP7,CER1,MAP3K7,VANGL2,WNT2B,WNT11,WNT10B,DKK2,LOC728622,CHP2,AXIN1,AXIN2,DKK4,NFAT5,MYC,SOX17,CSNK2A1,CSNK2A2,NFATC4,CSNK1A1,NFATC3,CSNK1E,BTRC,PRKX,SKP1,FBXW11,RBX1,CSNK2B,SIAH1,TBL1Y,WNT5B,CCND1,CAMK2A,NLK,CAMK2B,CAMK2D,CAMK2G,PRKACA,APC,PRKACB,PRKACG,WNT16,DAAM1,CHD8,FRAT1,CACYBP,CCND2,NFATC2,NFATC1,CCND3,PLCB2,PLCB1,CSNK1A1L,PRKCB,PLCB3,PRKCA,PLCB4,WIF1,PRICKLE2,PORCN,RHOA,FRAT2,PRKCG,MAPK9,MAPK10,WNT3A,DVL3,RAC2,DVL2,RAC3,FZD3,DKK1,CXXC4,DVL1,FOSL1,CUL1,WNT10A,WNT4,SMAD3,TCF7,SMAD4,RAC1,TCF7L2,SMAD2,WNT1,MAPK8,EP300,WNT7A,GSK3B,WNT7B,PSEN1,WNT8A,WNT8B,WNT2,WNT3,WNT5A,WNT6,CTNNB1,PPP2CB,PPP2CA,PPP2R1A,TBL1X,PPP2R1B,ROCK2,NKD1,FZD10,FZD5,NKD2,TCF7L1,RUVBL1,PPARD,PPP3CB,TP53,PPP3CA,PPP2R5A,PPP2R5E,PPP2R5D,PPP2R5C,PPP2R5B,FZD2,SFRP5,SFRP4,CHP |
| BIOCARTA WNT PATHWAY | CSNK2A1,WNT1,CSNK1D,CSNK1A1,GSK3B,BTRC,WIF1,CTNNB1,FZD1,PPP2CA,CREBBP,LEF1,TLE1,CCND1,CTBP1,NLK,APC,MAP3K7,DVL1,PPARD,FRAT1,AXIN1,HDAC1,TAB1,SMAD4,MYC, |
| KEGG HEDGEHOG SIGNALING PATHWAY | CSNK1A1L,HHIP,PTCH2,GAS1,WNT3A,ZIC2,WNT9B,WNT9A,LRP2,CSNK1G1,WNT2B,WNT11,WNT10B,IHH,SMO,WNT10A,WNT4,CSNK1G3,SHH,WNT1,CSNK1D,RAB23,CSNK1A1,CSNK1G2,CSNK1E,BMP8A,GSK3B,WNT7A,BTRC,WNT7B,WNT8A,WNT8B,WNT2,WNT3,PRKX,WNT5A,WNT6,FBXW11,STK36,WNT5B,GLI1,DHH,PRKACA,PRKACB,SUFU,BMP4,PRKACG,BMP2,GLI2,BMP7,GLI3,PTCH1,BMP8B,WNT16,BMP5,BMP6, |
| PID HIF1APATHWAY | HIF1A,OS9,VHL,HSP90AA1,HIF1AN,TCEB1,NAA10,TCEB2,COPS5,TP53,RBX1,ARNT,EGLN3,EGLN1,CUL2,CDKN2A,GNB2L1,HIF3A,EGLN2 |
| PID TGFBRPATHWAY | EIF2A,PARD6A,NEDD4L,CTNNB1,OCLN,SPTBN1,RPS6KB1,TAB2,TGFB3,CAV1,TGFB1,GRB2,RNF111,PPP2CA,PPP2CB,SMURF1,SMAD7,DAXX,WWP1,TGFB2,DAB2,PDPK1,SOS1,AXIN1,ITCH,RHOA,PPP2R2A,TGFBR1,SMURF2,YWHAE,ZFYVE16,PML,SMAD2,PPP1CA,DACT2,CTGF,STRAP,PPP1R15A,FKBP1A,ARRB2,SMAD4,CAMK2A,SMAD3,DYNLRB1,SKIL,SHC1,MAP3K7,TAB1,XIAP,BAMBI,TGFBRAP1,ZFYVE9,YAP1,TGFBR2,TGFBR3 |
| PID WNT SIGNALING PATHWAY | DKK1,WNT3A,FZD9,WNT2,WNT7A,IGFBP4,WNT1,FZD8,FZD1,KREMEN2,LRP6,KREMEN1,FZD2,LRP5,FZD6,RSPO1,WNT7B,FZD4,CTHRC1,WNT5A,RYK,FZD5,ATP6AP2,WNT3,ROR2,WIF1,FZD10,FZD7 |
| PID WNT CANONICAL PATHWAY | LRP6,GSK3A,AXIN1,CUL3,WNT3A,DVL3,GSK3B,FZD5,PPP2R5A,CSNK1G1,DVL1,PI4K2A,CTNNB1,DVL2,APC,NKD2,PIP5K1B,RANBP3,KLHL12,CAV1, |
| PID WNT NONCANONICAL PATHWAY | YES1,PRKCZ,MAP3K7,TAB2,FLNA,CSNK1A1,CDC42,DAAM1,FZD7,WNT5A,ROR2,NFATC2,SETDB1,NLK,ROCK1,RAC1,ARRB2,FZD5,MAPK8,CHD7,MAPK9,PPARG,CAMK2A,DVL1,RHOA,FZD6,CTHRC1,DVL2,DVL3,FZD2,TAB1,MAPK10, |
| PID HEDGEHOG GLIPATHWAY | SUFU,HDAC2,SSPO,PIAS1,GLI2,HDAC1,GNB1,SMO,RAB23,RBBP7,GLI3,CSNK1G3,KIF3A,CREBBP,GLI1,STK36,GNG2,GSK3B,FBXW11,GNAI3,SHH,CSNK1G2,XPO1,CSNK1E,IFT88,CSNK1D,MAP2K1,MTSS1,PRKCD,CSNK1A1,AKT1,GNAZ,IFT172,GNAI1,PTCH1,LGALS3,GNAO1,CSNK1G1,FOXA2,SIN3B,RBBP4,SIN3A,SPOP,ARRB2,GNAI2,SAP30,SAP18,PRKACA |

| | |
|---|---|
| KEGG JAK STAT SIGNALING PATHWAY | STAT3,STAT4,STAT1,STAT2,PIAS3,TYK2,IL21R,CREBBP,SOCS5,IL23A,CTF1,STAM,SPRY4,IL22,JAK1,AKT1,AKT2,JAK3,JAK2,AKT3,STAT5B,STAT5A,TPO,STAT6,MYC,IL24,SPRED1,PIK3R5,CSF2,IL13RA2,PIAS1,IL13RA1,CNTF,PRLR,CNTFR,PRL,SOCS1,IFNE,CSF2RB,CSF2RA,IL23R,CSF3,SOCS3,CCND1,IL22RA2,CSH1,BCL2L1,CSF3R,PIAS4,CBLC,CLCF1,OSMR,IL20,CCND2,CCND3,SOS2,EPO,PTPN11,IL9R,MPL,SPRED2,IFNA5,IFNA4,IFNA2,IL28RA,SOCS2,IFNA1,CISH,PTPN6,IL9,IL7R,IL12RB1,IL12B,IL13,IL12RB2,IL11RA,IL12A,IRF9,IFNA17,PIAS2,IFNA21,IFNA6,LEPR,IFNA7,IFNA8,IFNA10,IFNA13,GRB2,IFNA14,LEP,IFNA16,IL26,IL19,IL10,IL10RA,SOS1,IL10RB,EPOR,IL11,IL3RA,IL3,EP300,IL2RG,IL2RB,CBLB,OSM,CBL,IL15RA,IFNGR2,IL15,IFNG,IFNGR1,IFNB1,IFNAR1,IFNAR2,IL21,IL20RB,IL2RA,IL22RA1,IL20RA,IL2,IL6R,PIK3R3,CRLF2,IFNK,SPRY2,IL6ST,SPRY1,IL7,STAM2,GHR,IL28B,IL29,SOCS4,PIK3CA,PIK3CB,PIM1,IL28A,PIK3CD,LIF,GH1,GH2,IFNW1,LIFR,TSLP,IL4,IL4R,SPRY3,IL5,PIK3CG,SOCS7,IL5RA,PIK3R1,IL6,PIK3R2, |
| PID NOTCH PATHWAY | NCOR1,NOTCH1,DTX1,FURIN,SSPO,DLL1,DNM1,PTCRA,RAB11A,NEURL,CTBP1,FBXW7,NUMB,CBL,MFAP2,MYCBP,DLL4,MAML1,SPEN,MYC,SKP2,CNTN6,NOTCH2,PSEN1,MAML2,YY1,GATA3,ENO1,ITCH,IL4,APH1B,LNX1,NCOR2,RBPJ,CDKN1A,JAG2,ADAM12,DLK1,EPS15,DNER,SKP1,NOTCH3,RBBP8,NOTCH4,JAG1,APH1A,DLL3,PSENEN,MFAP5,NCSTN,CCND1,EP300,HDAC1,MARK2,MIB1,ADAM10,CUL1,CNTN1,KDM1A, |
| KEGG NOTCH SIGNALING PATHWAY | HES5,DTX3,NOTCH4,DTX3L,NOTCH3,NOTCH2,EP300,HES1,NOTCH1,NUMB,PSEN2,PSEN1,PTCRA,SNW1,APH1A,KAT2A,ADAM17,RFNG,RBPJ,DTX1,CREBBP,DTX2,MAML1,CTBP2,NCOR2,CTBP1,DVL3,JAG2,DVL2,NUMBL,MAML2,KAT2B,DLL4,PSENEN,DLL3,DVL1,CIR1,DLL1,LFNG,JAG1,MAML3,HDAC1,HDAC2,NCSTN,DTX4,MFNG,RBPJL, |
| REACTOME G1 S TRANSITION | CDK2,PSMD14,CDK7,CDKN1A,CDKN1B,DBF4,CKS1B,DHFR,E2F1,ORC6,ORC3,POLA2,FBXO5,PPP2R3B,RPA4,MAX,MCM2,MCM3,MCM4,MCM5,MCM6,MCM7,MNAT1,LOC441488,MYC,ORC1,ORC2,ORC4,ORC5,PCNA,POLA1,POLE,POLE2,PPP2CA,PPP2CB,PPP2R1A,PPP2R1B,MCM10,PRIM1,PRIM2,PSMA1,PSMA2,PSMA3,PSMA4,PSMA5,PSMA6,PSMA7,PSMB1,PSMB2,PSMB3,PSMB4,PSMB5,PSMB6,PSMB7,PSMB8,PSMB9,PSMB10,PSMC1,PSMC2,PSMC3,PSMC4,PSMC5,PSMC6,PSMD1,PSMD2,PSMD3,PSMD4,PSMD5,PSMD7,PSMD8,PSMD9,PSMD10,PSMD11,PSMD12,PSMD13,PSME1,PSME2,RB1,RPA1,RPA2,RPA3,RPS27A,RRM2,DHFRP1,LOC645084,SKP1,SKP2,LOC652826,TFDP1,TK2,RPS27AP11,TYMS,LOC729964,UBA52,WEE1,CDT1,CDC7,CDC45,MCM8,CUL1,CCNA2,CCNA1,CCNB1,CCNE1,CCNH,PKMYT1,CCNE2,PSMF1,CDK1,PSMD6,CDC6,CDC25A |
| REACTOME SIGNALING BY NOTCH | HDAC6,HDAC5,MAMLD1,ADAM10,CDK8,ST3GAL6,TMED2,DTX2,CNTN1,CREBBP,MIB2,JAG1,DTX1,E2F1,E2F3,EIF2C3,EIF2C4,EP300,SNW1,TNRC6B,DTX4,NCSTN,HEY1,HEY2,POFUT1,KAT2A,HEYL,EIF2C1,B4GALT1,EIF2C2,TNRC6A,DLL1,HDAC1,HDAC2,HIF1A,HES1,RBPJ,JAG2,JUN,HES5,LFNG,ARRB1,ARRB2,MFNG,MOV10,LOC441488,MYC,NOTCH2,NOTCH3,NOTCH4,ATP2A1,ATP2A2,ATP2A3,FURIN,APH1A,HDAC7,DLL4,FBXW7,MAML3,PSENEN,HDAC8,PSEN1,PSEN2,POGLUT1,MIB1,TNRC6C,RAB6A,CCND1,RFNG,RPS27A,SEL1L,ST3GAL3,SKP1,ADAM17,TBL1X,TFDP1,TLE1,TLE2,TLE3,TLE4,TP53,LOC728030,RPS27AP11,UBA52,TBL1XR1,HDAC11,APH1B,HDAC10,MAML2,CUL1,NUMB,DLK1,HDAC3,KAT2B,CCNC,NEURL,DNER,NCOR1,NCOR2,HDAC9,HDAC4,MAML1,RBX1 |
| REACTOME SIGNALING BY WNT | FRAT1,PSMD14,FAM123B,PSMA8,CSNK1A1,CTNNB1,PSME4,FRAT2,APC,PPP2CA,PPP2CB,PPP2R1A,PPP2R1B,PPP2R5A,PPP2R5B,PPP2R5C,PPP2R5D,PPP2R5E,PSMA1,PSMA2,PSMA3,PSMA4,PSMA5,PSMA6,PSMA7,PSMB1,PSMB2,PSMB3,PSMB4,PSMB5,PSMB6,PSMB7,PSMB8,PSMB9,PSMB10,PSMC1,PSMC2,PSMC3,PSMC4,PSMC5,PSMC6,PSMD1,PSMD2,PSMD3,PSMD4,PSMD5,PSMD7,PSMD8,PSMD9,PSMD10,PSMD11,PSMD12,PSMD13,PSME1,PSME2,RPS27A,SKP1,LOC652826,RPS27AP11,UBA52,AXIN1,CUL1,BTRC,PSMF1,PSMD6, |
| REACTOME DOWNREGULATION OF TGF BETA RECEPTOR SIGNALING | STUB1,STRAP,PPP1R15A,SMAD2,SMAD3,SMAD7,UCHL5,PPP1CA,PPP1CB,PPP1CC,PMEPA1,SMURF1,RPS27A,SMURF2,TGFB1,TGFBR1,TGFBR2,RPS27AP11,UBA52,LOC731429,XPO1,MTMR4,ZFYVE9, |
| REACTOME TGF BETA RECEPTOR SIGNALING IN EMT EPITHELIAL TO MESENCHYMAL TRANSITION | FKBP1A,ARHGEF18,RHOA,F11R,PARD6A,PRKCZ,PARD3,SMURF1,CGN,RPS27A,TGFB1,TGFBR1,TGFBR2,RPS27AP11,UBA52,LOC731429 |
| REACTOME TGF BETA RECEPTOR SIGNALING ACTIVATES SMADS | STUB1,STRAP,FKBP1A,PPP1R15A,SMAD2,SMAD3,SMAD4,SMAD7,FURIN,UCHL5,PPP1CA,PPP1CB,PPP1CC,PMEPA1,SMURF1,RPS27A,SMURF2,TGFB1,TGFBR1,TGFBR2,RPS27AP11,UBA52,LOC731429,XPO1,MTMR4,ZFYVE9, |

| | |
|---|---|
| REACTOME SIGNALING BY TGF BETA RECEPTOR COMPLEX | CDK8,CDK9,STUB1,CDKN2B,STRAP,PARP1,E2F4,E2F5,FKBP1A,ARHGEF18,PPP1R15A,WWTR1,HDAC1,JUNB,RHOA,SMAD2,SMAD3,SMAD4,SMAD7,MEN1,LOC441488,MYC,FURIN,SERPINE1,F11R,PARD6A,UCHL5,TRIM33,PPM1A,PPP1CA,PPP1CB,PPP1CC,PRKCZ,PARD3,PMEPA1,SMURF1,CGN,RBL1,TGIF2,RPS27A,SMURF2,SKI,SKIL,SP1,TFDP1,TGFB1,TGFBR1,TGFBR2,TGIF1,RPS27AP11,UBA52,LOC731429,UBE2D1,UBE2D3,XPO1,USP9X,CCNC,CCNT1,CCNT2,MTMR4,ZFYVE9,NCOR1,NCOR2 |
| REACTOME P53 INDEPENDENT G1 S DNA DAMAGE CHECKPOINT | PSMD14,CHEK1,CHEK2,PSMA8,PSME4,ATM,PSMA1,PSMA2,PSMA3,PSMA4,PSMA5,PSMA6,PSMA7,PSMB1,PSMB2,PSMB3,PSMB4,PSMB5,PSMB6,PSMB7,PSMB8,PSMB9,PSMB10,PSMC1,PSMC2,PSMC3,PSMC4,PSMC5,PSMC6,PSMD1,PSMD2,PSMD3,PSMD4,PSMD5,PSMD7,PSMD8,PSMD9,PSMD10,PSMD11,PSMD12,PSMD13,PSME1,PSME2,RPS27A,LOC651610,LOC652826,RPS27AP11,UBA52,PSMF1,PSMD6,CDC25A |
| REACTOME P53 DEPENDENT G1 DNA DAMAGE RESPONSE | CDK2,PSMD14,CDKN1A,CDKN1B,PSMA8,PSME4,MDM2,ATM,PSMA1,PSMA2,PSMA3,PSMA4,PSMA5,PSMA6,PSMA7,PSMB1,PSMB2,PSMB3,PSMB4,PSMB5,PSMB6,PSMB7,PSMB8,PSMB9,PSMB10,PSMC1,PSMC2,PSMC3,PSMC4,PSMC5,PSMC6,PSMD1,PSMD2,PSMD3,PSMD4,PSMD5,PSMD7,PSMD8,PSMD9,PSMD10,PSMD11,PSMD12,PSMD13,PSME1,PSME2,RPS27A,RFWD2,LOC651610,LOC652826,TP53,RPS27AP11,RFWD2P1,UBA52,CCNE1,CCNE2,PSMF1,PSMD6 |
| REACTOME INHIBITION OF REPLICATION INITIATION OF DAMAGED DNA BY RB1 E2F1 | E2F1,POLA2,PPP2R3B,LOC441488,POLA1,PPP2CA,PPP2CB,PPP2R1A,PPP2R1B,PRIM1,PRIM2,RB1,TFDP1 |
| REACTOME G2 M DNA DAMAGE CHECKPOINT | CHEK1,CHEK2,ATM,ATR,LOC648152,LOC651610,LOC651921,WEE1,ATRIP,CCNB1,CDK1,CDC25C |
| REACTOME DNA REPAIR | RAD50,CDK7,CCNO,MAD2L2,POLD3,ERCC8,ALKBH2,DDB1,DDB2,ERCC1,ERCC2,ERCC3,ERCC4,ERCC5,ERCC6,FANCA,FANCC,FANCD2,FANCE,FANCB,FANCF,FANCG,ALKBH3,FEN1,SMUG1,XRCC6,ZBTB32,UBE2T,GTF2H1,GTF2H2,GTF2H3,GTF2H4,H2AFX,APEX1,LOC389901,LIG1,LIG3,LIG4,MGMT,MNAT1,MPG,MRE11A,MUTYH,NBN,ATM,NTHL1,OGG1,PCNA,REV1,POLB,POLD1,POLD2,POLE,POLE2,POLH,POLR2A,POLR2B,POLR2C,POLR2D,POLR2E,POLR2F,POLR2G,POLR2H,POLR2I,POLR2J,POLR2K,POLR2L,ATR,FANCL,TDP1,PRKDC,XAB2,FANCM,POLD4,RAD23B,RAD51,RAD52,REV3L,RFC2,RFC3,RFC4,RFC5,RPA1,RPA2,RPA3,RPS27A,LOC648152,LOC651610,LOC651921,LOC652672,LOC652857,GTF2H2B,BRCA1,BRCA2,TCEA1,TDG,TP53BP1,RPS27AP11,UBA52,USP1,XPA,XPC,XRCC1,XRCC4,XRCC5,PALB2,C17orf70,BRIP1,MBD4,CCNH,C19orf40,MDC1 |
| REACTOME REGULATION OF APOPTOSIS | PSMD14,PSMA8,DAPK1,DAPK3,DCC,UNC5B,PSME4,DAPK2,APPL1,PAK2,PSMA1,PSMA2,PSMA3,PSMA4,PSMA5,PSMA6,PSMA7,PSMB1,PSMB2,PSMB3,PSMB4,PSMB5,PSMB6,PSMB7,PSMB8,PSMB9,PSMB10,PSMC1,PSMC2,PSMC3,PSMC4,PSMC5,PSMC6,PSMD1,PSMD2,PSMD3,PSMD4,PSMD5,PSMD7,PSMD8,PSMD9,PSMD10,PSMD11,PSMD12,PSMD13,PSME1,PSME2,RPS27A,LOC652826,RPS27AP11,UBA52,ARHGAP10,CASP3,CASP9,UNC5A,PSMF1,MAGED1,PSMD6 |
| REACTOME APOPTOSIS | BCL2L11,DNM1L,BCAP31,PSMD14,ADD1,DYNLL2,PSMA8,CTNNB1,DAPK1,DAPK3,DCC,DFFA,DFFB,DSG1,DSG2,DSG3,DSP,E2F1,AKT1,UNC5B,ACIN1,PSME4,FNTA,DAPK2,APPL1,GAS2,BBC3,DBNL,GSN,GZMB,H1F0,HIST1H1C,HIST1H1D,HIST1H1E,HIST1H1B,HIST1H1A,HMGB1,HMGB2,APAF1,APC,BIRC2,XIAP,FAS,FASLG,KPNA1,KPNB1,LMNA,LMNB1,MAPT,LOC441488,NMT1,OCLN,PAK2,MST4,PKP1,PLEC,PMAIP1,CYCS,PPP3R1,PRKCD,PRKCQ,MAPK8,DIABLO,PSMA1,PSMA2,PSMA3,PSMA4,PSMA5,PSMA6,PSMA7,PSMB1,PSMB2,PSMB3,PSMB4,PSMB5,PSMB6,PSMB7,PSMB8,PSMB9,PSMB10,PSMC1,PSMC2,PSMC3,PSMC4,PSMC5,PSMC6,PSMD1,PSMD2,PSMD3,PSMD4,PSMD5,PSMD7,PSMD8,PSMD9,PSMD10,PSMD11,PSMD12,PSMD13,BAD,PSME1,PSME2,PTK2,BAK1,BAX,BCL2,BCL2L1,ROCK1,RPS27A,SATB1,BID,LOC647859,LOC652460,LOC652826,BMX,SPTAN1,TFDP1,TJP1,TNF,TNFRSF1A,TP53,TRAF2,ROCK1P1,RPS27AP11,UBA52,VIM,YWHAB,ARHGAP10,CASP3,CASP6,CASP7,CASP8,CASP9,STK24,CASP10,DYNLL1,TRADD,RIPK1,TNFSF10,FADD,TNFRSF10B,CFLAR,UNC5A,BMF,TJP2,PSMF1,MAGED1,PSMD6,CDH1 |
| REACTOME INTRINSIC PATHWAY FOR APOPTOSIS | BCL2L11,DYNLL2,E2F1,AKT1,BBC3,GZMB,APAF1,XIAP,LOC441488,NMT1,PMAIP1,CYCS,PPP3R1,MAPK8,DIABLO,BAD,BAK1,BAX,BCL2,BCL2L1,BID,TFDP1,TP53,YWHAB,CASP3,CASP7,CASP8,CASP9,DYNLL1,BMF |

| | |
|---|---|
| KEGG APOPTOSIS | CASP10,CASP9,CASP8,CASP7,CHUK,PRKAR2B,TNF,TNFSF10,BIRC3,XIAP,PPP3R2,PPP3CC,PPP3R1,MYD88,FADD,CFLAR,RIPK1,BAD,IRAK4,BID,BAX,IKBKB,CASP6,IL1A,AKT1,CASP3,AKT2,TNFRSF1A,AKT3,CHP2,ATM,ENDOG,NFKB1,NFKBIA,CAPN2,PIK3R5,IKBKG,CAPN1,IL3RA,IL3,RELA,ENDOD1,APAF1,PRKX,CSF2RB,TNFRSF10A,TRAF2,TNFRSF10D,NGF,TNFRSF10B,TNFRSF10C,MAP3K14,IL1RAP,IL1B,IRAK2,IL1R1,IRAK1,TRADD,PIK3R3,BCL2,BCL2L1,BIRC2,IRAK3,PRKACA,PRKACB,PRKACG,PPP3CB,TP53,PPP3CA,PIK3CA,PIK3CB,FAS,DFFA,CYCS,DFFB,PIK3CD,LOC651610,PRKAR1A,FASLG,PRKAR2A,PRKAR1B,EXOG,PIK3CG,AIFM1,NTRK1,PIK3R1,PIK3R2,CHP |
| REACTOME PI3K CASCADE | FRS2,THEM4,KLB,DOK1,EIF4B,EIF4E,EIF4EBP1,EIF4G1,AKT2,FGF1,FGF2,FGF3,FGF4,FGF5,FGF6,FGF7,FGF8,FGF9,FGF10,FGFR1,FGFR3,FGFR2,FGFR4,MTOR,GAB1,FGF20,FGF22,GRB2,EEF2K,PIK3R4,INS,INSR,IRS1,PDE3B,PRKAG2,PDPK1,CAB39,PIK3C3,PIK3CA,PIK3CB,PIK3R1,PIK3R2,PRKAG3,TLR9,PPM1A,STRADB,PRKAA1,PRKAA2,PRKAB1,PRKAB2,PRKAG1,RPTOR,TRIB3,RHEB,RPS6,RPS6KB1,MLST8,LOC644462,STK11,TSC1,TSC2,LOC729120,LOC730244,FGF23,CAB39L,IRS2,FGF18,FGF17,STRADA,KL,FGF19, |
| REACTOME PI3K EVENTS IN ERBB4 SIGNALING | AKT3,CDKN1A,CDKN1B,CHUK,THEM4,CREB1,NRG4,HBEGF,ERBB4,EREG,AKT1,AKT2,FOXO1,FOXO3,PHLPP1,MTOR,RICTOR,GSK3A,NRG1,NR4A1,MDM2,FOXO4,PDPK1,PIK3CA,PIK3R1,BAD,PTEN,TRIB3,RPS6KB2,MLST8,BTC,TSC2,LOC729120,LOC731292,MAPKAP1,CASP9,AKT1S1,NRG2, |
| REACTOME PI3K EVENTS IN ERBB2 SIGNALING | AKT3,CDKN1A,CDKN1B,CHUK,THEM4,CREB1,NRG4,HBEGF,EGF,EGFR,ERBB2,ERBB3,ERBB4,EREG,AKT1,AKT2,FOXO1,FOXO3,PHLPP1,MTOR,RICTOR,GAB1,GRB2,GSK3A,NRG1,NR4A1,MDM2,FOXO4,PDPK1,PIK3CA,PIK3R1,BAD,PTEN,TRIB3,RPS6KB2,MLST8,BTC,TSC2,LOC729120,LOC731292,MAPKAP1,CASP9,AKT1S1,NRG2, |
| BIOCARTA PTEN PATHWAY | PTK2,PTEN,PDK2,SHC1,FOXO3,PDPK1,BCAR1,CDKN1B,ILK,AKT1,PIK3CA,ITGB1,FASLG,GRB2,MAPK3,PIK3R1,SOS1,MAPK1 |
| REACTOME NEGATIVE REGULATION OF THE PI3K AKT NETWORK | AKT3,THEM4,AKT1,AKT2,PHLPP1,PTEN,TRIB3,LOC729120,LOC731292 |
| REACTOME PI3K AKT ACTIVATION | AKT3,CDKN1A,CDKN1B,CHUK,THEM4,CREB1,AKT1,AKT2,FOXO1,FOXO3,PHLPP1,MTOR,RICTOR,GSK3A,NR4A1,IRS1,RHOA,MDM2,FOXO4,NGF,NTRK1,PDPK1,PIK3CA,PIK3CB,PIK3R1,PIK3R2,BAD,PTEN,TRIB3,RPS6KB2,MLST8,TSC2,LOC729120,LOC731292,MAPKAP1,CASP9,AKT1S1,IRS2, |
| REACTOME SIGNALLING TO RAS | MAPK14,SHC2,GRB2,HRAS,KRAS,NGF,NRAS,NTRK1,SHC3,MAPK1,MAPK3,MAPK11,MAPK13,MAP2K1,MAP2K2,RAF1,RALA,RALB,RALGDS,MAPK12,SHC1,SOS1,SRC,YWHAB,MAPKAPK3,MAPKAPK2,CDK1 |
| BIOCARTA RAS PATHWAY | CASP9,HRAS,FOXO4,CHUK,RALA,RAF1,BCL2L1,ELK1,RELA,MAP2K1,RALBP1,RALGDS,AKT1,PIK3CA,RHOA,PLD1,MAPK3,BAD,NFKB1,PIK3CG,RAC1,PIK3R1,CDC42, |

| | |
|---|---|
| KEGG MAPK SIGNALING PATHWAY | JUN,MEF2C,ELK4,ELK1,JUND,GADD45B,ZAK,STMN1,RRAS2,MAP3K5,MAP3K1,MAP3K3,MAP3K4,MAP3K7,MAP3K8,AKT1,AKT2,ARRB2,CD14,ARRB1,NRAS,DUSP16,CHP2,RASGRP3,NFKB2,NFKB1,MYC,NFATC4,MAPK14,FLNC,FLNA,KRAS,FLNB,PRKX,TRAF6,TGFB2,DUSP1,DUSP2,TGFB1,TRAF2,BDNF,TAB2,ECSIT,TGFBR2,DUSP7,TGFBR1,DUSP5,DUSP6,DUSP3,TGFB3,PLA2G4E,DUSP4,CACNG5,CACNG4,NF1,PLA2G12A,NFATC2,RASGRP4,MAP3K2,MAX,DUSP10,FGF9,FGF8,FGF7,FGF6,FGF5,FGF3,FGF4,FGF1,FGF2,PTPN5,FGF21,IL1R2,MAPK9,CACNA2D3,MAPK10,MAPK11,RASGRP2,PLA2G2A,MAP2K2,PLA2G4A,MAP2K3,MECOM,PLA2G5,MAPK13,MAP2K1,FGFR2,MAP2K7,RASA2,MAPK8IP2,RASGRF1,MAP2K5,RASGRF2,FGFR4,MAP2K6,MAPK8IP3,MAP3K6,CASP3,MAP3K12,FGFR3,FGFR1,RASA1,FGF14,RPS6KA2,RPS6KA3,FGF17,FGF16,FGF10,GRB2,FGF11,FGF12,FGF13,PLA2G1B,RPS6KA1,MAPKAPK3,IKBKG,HRAS,CACNG2,FGF23,CACNG3,MKNK2,FGF18,STK4,STK3,MAPK8IP1,MOS,RAP1A,MAPT,RAP1B,PPP3CB,PPP3CA,CACNA1H,CACNA1G,CACNA1I,ATF4,TAB1,FOS,TAOK2,RPS6KA6,PPP3R2,CACNG8,PPP5C,PPP3CC,CACNG6,PPP3R1,CACNG7,MAP2K4,ATF2,PDGFRB, JMJD7-PLA2G4B,MAP4K3,PLA2G6,PLA2G2E,PLA2G10,MAP4K4,RPS6KA5,BRAF,IKBKB,PLA2G4B,MAP3K11,CACNA2D2,IL1A,PLA2G2F,DAXX,AKT3,GADD45G,FGF20,RELB,MAPKAPK5,MAPK12,RELA,GNA12,HSPA8,HSPB1,PTPRR,LAMTOR3,GADD45A,NGF,DDIT3,MAP3K14,TAOK1,PDGFA,FGF22,PDGFB,NLK,PDGFRA,PRKACA,PRKACB,PAK1,PRKACG,CRK,CDC25B,CRKL,MAP3K13,PLA2G2D,CDC42,RASGRP1,CACNA2D1,CACNB1,SRF,CACNB2,SOS2,CACNB3,CACNB4,CHUK,CACNG1,PRKCB,RAF1,PRKCA,TNF,PAK2,MKNK1,PLA2G3,PRKCG,PTPN7,RAPGEF2,HSPA1L,CACNA1A,HSPA1B,RAC2,HSPA2,CACNA1D,CACNA1E,RAC3,CACNA1B,HSPA1A,PLA2G12B,CACNA1C,DUSP8,CACNA1F,CACNA1S,MAP4K1,TNFRSF1A,DUSP9,PPM1A,PPM1B,MAPK3,RPS6KA4,CACNA2D4,HSPA6,MAPK7,FGF19,RAC1,SOS1,MAPK1,DUSP14,MAP4K2,MAPK8,EGFR,MAPKAPK2,EGF,RRAS,TAOK3,GNG12,NTF4,IL1B,MRAS,NTF3,IL1R1,TP53,FAS,NR4A1,PLA2G2C,FASLG,NTRK2,NTRK1,CHP |
| BIOCARTA MAPK PATHWAY | MEF2C,JUN,MEF2D,MAX,MEF2BNB-MEF2B,CHUK,RAF1,MEF2A,STAT1,SHC1,ELK1,PAK2,CEBPA,MKNK1,MAP2K4,RIPK1,ATF2,RAPGEF2,MAPK9,MAP3K5,MAP4K3,MAPK10,MAPK11,MAP2K2,MAP4K4,BRAF,RPS6KA5,MAP2K3,MAP3K1,MAPK13,MAP3K3,MAP2K1,MAP3K4,MAP2K7,MAP3K10,MAP3K9,IKBKB,MAP2K5,MAP3K11,CREB1,MAP2K6,MAP3K7,MAP3K8,MAP3K12,MAP3K6,MAP4K1,MAP4K5,RPS6KA2,RPS6KA3,DAXX,RPS6KB1,RPS6KB2,GRB2,MAPK3,MAPK4,RPS6KA4,MAPK6,NFKB1,MAPK7,RAC1,NFKBIA,MYC,RPS6KA1,MAPK1,MAPKAPK3,HRAS,MAP4K2,MAPK8,MAPKAPK5,MAPK14,MAPK12,RELA,MAPKAPK2,TGFB2,MKNK2,TGFB1,TRAF2,ARAF,MAP3K14,TGFBR1,TRADD,TGFB3,PAK1,SP1,MAP3K13,FOS,MAP3K2 |
| KEGG CELL CYCLE | CDC16,CDC7,CDC45,GADD45B,DBF4,ANAPC1,CREBBP,MDM2,ABL1,SMC1B,LOC728622,GADD45G,ATM,ATR,ANAPC7,RBL2,ANAPC5,RBL1,MYC,CDC14B,SMC1A,CDC14A,LOC731751,SKP1,TGFB2,TGFB1,GADD45A,STAG1,LOC650621,PLK1,RBX1,STAG2,TGFB3,MCM4,ORC6,CCND1,MAD1L1,MCM3,MCM6,MCM5,YWHAB,CCNA2,MCM7,BUB1,CHEK1,WEE2,MCM2,PTTG1,CDC27,CDC25B,CDC25C,LOC651610,CDC25A,CDC6,CDC20,BUB3,YWHAZ,CCND2,YWHAH,CCNB1,YWHAG,YWHAE,CCNE1,ORC3,CCND3,PTTG2,SFN,E2F1,TFDP1,ZBTB17,CDK1,ESPL1,ANAPC10,RAD21,BUB1B,ANAPC11,RB1,SKP2,CUL1,SMAD3,SMAD4,ANAPC2,TFDP2,PRKDC,MAD2L1,ANAPC4,SMAD2,YWHAQ,CHEK2,CDC23,EP300,GSK3B,CDKN2A,CDKN1C,CDKN1B,CDKN1A,CDKN2D,CCNA1,CDKN2B,CDKN2C,FZR1,SMC3,ANAPC13,PCNA,TTK,PKMYT1,CDK2,CDC26,E2F5,CDK4,WEE1,E2F4,E2F3,TP53,E2F2,ORC1,ORC2,CCNE2,CDK6,ORC4,CCNB2,CDK7,MAD2L2,ORC5,HDAC1,HDAC2,CCNH,CCNB3, |
| MIPS SWI SNF CHROMATIN REMODELING RELATED BRCA1 COMPLEX | ACTL6A,BRCA1,SMARCA2,SMARCA4,SMARCB1,SMARCC1,SMARCC2,SMARCD2,SMARCE1,ARID1A,ARID1B, |
| MIPS INO80 CHROMATIN REMODELING COMPLEX | ACTL6A,ACTR5,ACTR8,INO80C,INO80E,INO80D,INO80,MCRS1,NFRKB,RUVBL1,RUVBL2,TCF3,INO80B |
| MIPS SRCAP ASSOCIATED CHROMATIN REMODELING COMPLEX | ACTL6A,ACTR6,EAF2,H2AFZ,RUVBL1,RUVBL2,SRCAP,VPS72,YEATS4,ZNHIT1 |
| MIPS RNA POLYMERASE II COMPLEX CHROMATIN STRUCTURE MODIFYING | CCNC,CDK8,DRAP1,GTF2B,GTF2E1,GTF2F1,GTF2H1,MED21,PCSK4,POLR2A,SMARCB1,SMARCC1,SMARCC2,TBP,SMARCA2,SMARCA4,SMARCD1,SMARCD2,SMARCD3 |
| MIPS RNA POLYMERASE II COMPLEX CHROMATIN STRUCTURE MODIFYING 1 | ACTL6A,CCNC,CDK8,MED21,SMARCB1,SMARCC1,SMARCC2,SMARCE1,SMARCD1,SMARCD2,SMARCD3, |

| | |
|---|---|
| MIPS RNA POLYMERASE II COMPLEX CHROMATIN STRUCTURE MODIFYING 2 | CCNC,CDK8,CREBBP,ERCC3,GTF2B,GTF2F1,GTF2H3,MED21,KAT2B,POLR2A,SMARCA2,SMARCA4,SMARCB1 |
| KEGG AMINOACYL TRNA BIOSYNTHESIS | CARS2,DARS2,RARS,SARS,VARS2,YARS2,WARS,AARS,FARSA,HARS,SARS2,RARS2,NARS,LARS2,FARSB,AARS2,YARS,NARS2,GARS,IARS2,KARS,WARS2,TARSL2,LARS,VARS,PARS2,MARS2,CARS,IARS,TARS2,SEPSECS,MTFMT,TARS,DARS,HARS2,PSTK,QARS,EARS2,EPRS,FARS2,MARS, |
| PID NFKAPPABATYPICALPATHWAY | PIK3CA,NFKB1,CSNK2B,SYK,CSNK2A1,PIK3R1,SSPO,MAPK14,LCK,NFKBIA,RELA,SRC,BCL3,IKBKB,ARRB2,REL,CSNK2A2, |
| PID NFKAPPABCANONICALPATHWAY | CYLD,UBE2D3,TNFAIP3,NFKBIA,XPO1,MALT1,BIRC2,TRAF6,NOD2,PRKCA,RELA,RAN,TNF,IKBKG,NFKB1,SSPO,IKBKB,RIPK2,BCL10,CHUK,ATM,ERC1,TNFRSF1A, |
| MIPS TNF ALPHA NF KAPPA B SIGNALING COMPLEX | CHUK,IKBKG,KPNA3,NFKB1,NFKB2,NFKBIA,NFKBIB,NFKBIE,REL,RELA,RELB,TNIP2 |
| MIPS TNF ALPHA NF KAPPA B SIGNALING COMPLEX 1 | CHUK,FBXW7,IKBKB,IKBKG,MAP3K14,NFKB2,REL,RELA,SEC16A,USP2 |
| MIPS TNF ALPHA NF KAPPA B SIGNALING COMPLEX 2 | ANKRD28,BTRC,CHUK,CUL1,IKBKE,NFKB2,PPP6C,REL,RELA,PPP6R1,PPP6R2,SKP1 |
| MIPS TNF ALPHA NF KAPPA B SIGNALING COMPLEX 3 | CHUK,DDX3X,GLG1,GTF2I,IKBKG,MAP3K8,NFKB1,NFKB2,NFKBIA,NFKBIB,REL,RELA,RELB,RPL30,RPL6,RPS13,TNIP2, |
| MIPS TNF ALPHA NF KAPPA B SIGNALING COMPLEX 5 | CD3EAP,CHUK,CUL1,FBXW11,IKBKB,IKBKG,IQGAP2,KPNA2,LRPPRC,MCC,MTIF2,NFKB1,NFKB2,NFKBIB,PDCD2,POLR1A,POLR1B,POLR1D,POLR1E,POLR2H,POLR2L,RASAL2,REL,RELA,SKP1 |
| MIPS TNF ALPHA NF KAPPA B SIGNALING COMPLEX 6 | CDC37,CHUK,FBL,HSP90AA1,HSP90AB1,IKBKB,IKBKG,MAP3K14,RPL30,RPL4,RPL6,RPL8,RPS11,RPS13, |
| MIPS TNF ALPHA NF KAPPA B SIGNALING COMPLEX 10 | ATG16L1,CCAR1,CDC37,CHUK,HSP90AA1,HSP90AB1,IKBKG,NFKBIB,TBK1,TXLNA |
| PID P53REGULATIONPATHWAY | NEDD8,PPM1D,HIPK2,CHEK2,TP53,TP53AIP1,TRIM28,CCNG1,HUWE1,CSNK1E,SETD8,SKP2,ATM,CSE1L,CSNK1D,CSNK1A1,PPP2CA,CDK2,PIN1,CHEK1,SETD7,MDM4,KAT5,CCNA2,SMYD2,E4F1,EP300,PRMT5,KAT2B,PPP1R13L,CSNK1G1,RFWD2,KAT8,DAXX,DYRK2,TTC5,RPL5,RPL11,PRKCD,PPP2R4,MDM2,FBXO11,AKT1,CDKN2A,ATR,ABL1,CSNK1G2,GSK3B,CREBBP,RASSF1,MAPK14,MAPK9,RPL23,USP7,RCHY1,UBE2D1,CSNK1G3,YY1,MAPK8, |
| KEGG P53 SIGNALING PATHWAY | CASP9,CASP8,SFN,TSC2,IGF1,CDK1,GADD45B,RCHY1,ZMAT3,IGFBP3,RRM2B,BAI1,TP73,SERPINB5,RPRM,PPM1D,BID,MDM4,BAX,MDM2,TP53AIP1,CASP3,TP53I3,PMAIP1,PIDD,RFWD2,PERP,GADD45G,EI24,SESN2,ATM,ATR,SESN3,CHEK2,APAF1,CDKN2A,RRM2,CDKN1A,BBC3,GADD45A,TNFRSF10B,DDB2,GTSE1,SIAH1,CCND1,CD82,PTEN,CDK2,CHEK1,SESN1,CDK4,SERPINE1,TP53,CCNE2,FAS,CDK6,CYCS,STEAP3,CCNB2,LOC651610,CCND2,SHISA5,CCNB1,CCNG1,CCNE1,CCNB3,THBS1,CCNG2,CCND3 |
| BIOCARTA P53HYPOXIA PATHWAY | MAPK8,CSNK1D,ABCB1,CSNK1A1,EP300,HSPA1A,BAX,DNAJB1P1,HIF1A,HIC1,MDM2,TAF1,CDKN1A,AKT1,TP53,NQO1,FHL2,GADD45A,HSP90AA1,IGFBP3,ATM,RPA1,NFKBIB, |
| BIOCARTA P53 PATHWAY | CCND1,BCL2,E2F1,APAF1,BAX,CDK2,CDK4,MDM2,RB1,CDKN1A,TP53,GADD45A,ATM,TIMP3,PCNA,CCNE1 |
