## Supplementary material for "Drug-driven reclassification of multiple tumour subtypes reveals intrinsic molecular concordance of therapy across histologically disparate cancers": Supple Table 2

Supplementary Table 2. The enrichment of histological cancers in the most sensitive pharmacological subtype of PD0325901.

| Drug name | Drug ID | Target | Pathway | Cancer Type | pValue |
|---|---|---|---|---|---|
| PD0325901 | 1060 | MEK1, MEK2 | ERK MAPK signaling | COREAD | 4.58E-04 |
| | | | | HNSC | 3.59E-03 |
| | | | | PAAD | 4.17E-03 |
| | | | | SKCM | 1.30E-14 |
| | | | | THCA | 8.79E-04 |
